# Autoinhibition in the Signal Transducer CIN85 Modulates B Cell Activation

**DOI:** 10.1101/2023.07.31.551229

**Authors:** Daniel Sieme, Michael Engelke, Nasrollah Rezaei-Ghaleh, Stefan Becker, Jürgen Wienands, Christian Griesinger

**Affiliations:** Department for NMR-based Structural Biology, Max Planck Institute for Multidisciplinary Sciences, Am Fassberg 11, 37077 Göttingen, Germany; Institute for Cellular and Molecular Immunology, Georg-August University Göttingen, Humboldtallee 34, 37073 Göttingen, Germany; Institute of Physical Biology, Heinrich Heine University Düsseldorf, Universitätsstraße 1, 40225 Düsseldorf, Germany; Institute of Biological Information Processing, IBI-7: Structural Biochemistry, Forschungszentrum Jülich, Wilhelm-Johnen-Straße, 52428 Jülich, Germany

## Abstract

Signal transduction by the ligated B cell antigen receptor (BCR) depends on the pre-organization of its intracellular components, such as the effector proteins SLP65 and CIN85 within phase-separated condensates. These liquid-like condensates are based on the interaction between three Src homology 3 (SH3) domains and corresponding proline-rich recognition motifs (PRM) in CIN85 and SLP65, respectively. However, detailed information on the protein conformation and how it impacts on the capability of SLP65/CIN85 condensates to orchestrate BCR signal transduction is still lacking. This study identifies a hitherto unknown intramolecular SH3:PRM interaction between the C-terminal SH3 domain (SH3C) of CIN85 and an adjacent PRM. We used high-resolution nuclear magnetic resonance (NMR) experiments to study the flexible linker region containing the PRM and determined the extent of the interaction in multidomain constructs of the protein. Moreover, we observed that the phosphorylation of a serine residue located in the immediate vicinity of the PRM regulates this intramolecular interaction. This allows for a dynamic modulation of CIN85’s valency towards SLP65, regulating the extent of liquid-liquid phase separation. B cell culture experiments further revealed that the PRM/SH3C interaction is crucial for maintaining the physiological level of SLP65/CIN85 condensate formation, activation-induced membrane recruitment of CIN85, and subsequent mobilization of Ca^2+^. Our findings therefore suggest that the intramolecular interaction to the adjacent disordered linker is effective in modulating CIN85’s valency both *in vitro* and *in vivo*. This therefore constitutes a powerful way for the modulation of SLP65/CIN85 condensate formation and subsequent B cell signaling processes within the cell.

## Introduction

Scaffold proteins play an important role in the spatial and temporal organization of cellular processes and thus their significance for many of the interconnected signaling pathways cannot be overstated. Their efficient use of multiple modular domains and intrinsically disordered regions (IDR) enables the formation of the large macromolecular assemblies that play an important role in nearly all of the signaling pathways known within the cell ^1^. Specific functions range from recruiting effectors to specific subcellular locations ^2^, providing docking sites for the assembly of higher-order macromolecular structures ^3-4^ and to fine-tune the often weak and transient interactions within these assemblies ^5^. In particular, the combination of modular folded domains connected via IDR leads to multi-domain proteins with a large potential for internal dynamics, necessary for their many different functions. Consistently, the same scaffold protein can play different roles in separate signaling pathways, depending on differential splicing, post-translational modifications (PTM) and/or the presence of different other effector and scaffold proteins ^6^.

Cbl-interacting protein of 85 kDa (CIN85) is a protein expressed in many different cell types, involved in processes as diverse as cytokinesis ^7^, lysosomal degradation of the epidermal growth factor receptor ^8-9^, clathrin-mediated receptor internalization ^10^, cell adhesion and cytoskeletal remodeling ^11-12^, and both T cell receptor ^13^ and B cell receptor (BCR) signaling ^14-18^. In the context of processes associated with BCR signaling, CIN85 was shown to be constitutively associated with Src homology 2 domain-containing leukocyte protein of 65 kDa (SLP65) ^15^, engaging in promiscuous multivalent interactions between its Src-homology 3 (SH3) domains and SLP65 proline-rich motifs (PRMs) ^18^. By association with small unilamellar phospholipid vesicles via the N-terminal domain of SLP65, all these transient interactions lead to the formation of droplets showing characteristics of liquid-liquid phase separation (LLPS) ^17-18^. These organelles form in the resting state of the B cell and provide preformed complexes of CIN85 and SLP65, which allow for an accelerated cellular response upon BCR engagement ^17^. Notably, CIN85 is involved in the pre-assembly of effectors in resting T cells as well, but this involves a distinctly different set of interaction partners compared to B cells ^13^. Besides being necessary for the formation of droplets together with SLP65 in the resting state of the B cell, CIN85 itself also promotes higher-order structures: its C-terminal coiled-coil domain exhibits a high propensity for trimerization ^17^. In addition to the heterotypic interactions, PRMs inside the protein can compete with other motifs for binding to the SH3 domains ^6, 19^. In summary, the multitude of possible interactions leads to a complex network of transient interactions characteristic of a protein serving different contextual functions. However, how the cellular context leads to differential behavior of multi-domain proteins in distinct signaling pathways is still poorly understood. Dynamic regulation of these proteins is often facilitated by PTMs, such as phosphorylation at Tyr, Ser or Thr residues ^20^. Consequently, a common mode of regulation for multidomain proteins containing flexible linker regions is auto-inhibition ^21^. This is commonly caused by recognition motifs inside flexible linker or tail regions, occupying one of the domains completely until the interaction is perturbed. This results in the domain being able to engage with other effectors and/or become catalytically active ^22^. There is evidence by previous studies that CIN85 SH3 domains are able to recognize PRMs within disordered regions of CIN85, leading to intra-or intermolecular autoinhibition ^6, 19, 23-24^. Notably, Li *et al*.^*19*^ have provided indirect evidence of an autoinhibitory interaction mediated by the SH3C domain to the adjacent linker based on isothermal titration calorimetry experiments but did not investigate this further. Since the propensity for LLPS is mainly driven by the interaction of CIN85 SH3 domains with SLP65 PRMs, this mechanism could be a powerful way for the intracellular modulation of LLPS. These interactions have, however, not been characterized in detail in the context of the multidomain protein constructs, leaving the extent of their overall contributions to CIN85 protein conformation and function an open question.

In this study, we identified a hitherto unknown PRM that predominantly interacts with the CIN85 SH3C domain. Using NMR spectroscopy we characterized the interaction of the CIN85 SH3 domains with synthetic peptides, also addressing the influence of mutations on the binding. We assigned the backbone resonances of the disordered linker containing the motif and investigated SH3:PRM binding via NMR relaxation and translational diffusion experiments in multidomain protein constructs of various lengths. We determined that this SH3:PRM interaction modulated the valency of the CIN85 protein and therefore the extent of interaction with its constitutive binding partner SLP65. Finally, we showed the relevance of this interaction in DG75 B cell lymphoma cells for modulating B cell responses to stimulation of the BCR.

## Results

### The second IDR in CIN85 contains a novel PRM that in-teracts preferably with SH3C

We first investigated whether the linker regions in CIN85 were predicted to show deviations from a purely disordered linker. For this, we employed two predictors, that are based on flexible regions in high-resolution XRAY structures (DISOPRED3 ^25^) and backbone flexibility from NMR chemical shifts of IDPs (DynaMine ^26^). DISOPRED3 scores will be high for highly disordered sequences, while the DynaMine order parameter prediction indicates more rigid structures at high values. As displayed in Figure 1A, both predictors were able to distinguish the folded domains (SH3A-C and the coiled-coil (CC) domain) from the disordered linkers. In addition, both predictors show a significant deviation from a purely disordered sequence in the linker region between SH3B and SH3C (residues 162-263). Conserved residues in protein sequences can indicate functional importance, even in intrinsically disordered regions that typically do not show a high degree of conservation ^27^. We therefore determined the sequence conservation within the intrinsically disordered linker between SH3B and SH3C by performing a BLAST ^28^ search starting from the CIN85 Uniprot entry Q96B97-1 and compared the sequence conservation between different CIN85 homologs (Figure 1B). Inside the region predicted by DynaMine to show the lowest flexibility, we identified a novel proline-rich sequence (residues 223-230) that showed exceptional sequence conservation among all homologs tested.

**Figure 1:**
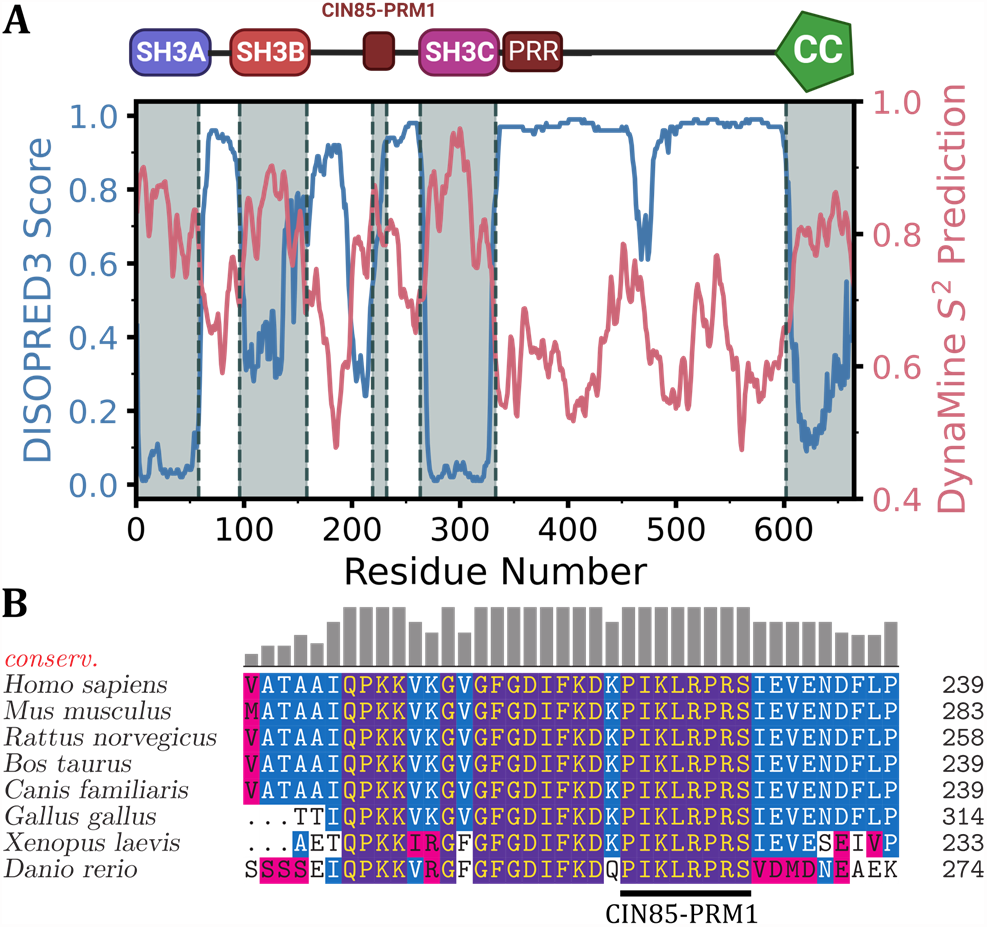
Sequence-based analysis of CIN85 linker regions. **A:** Prediction of disorder (DISOPRED3 ^25^) and flexibility (DynaMine ^26^) along the CIN85 amino acid sequence. DISOPRED3 scores will be close to one for highly disordered sequences, while the DynaMine order parameter pre-diction indicates more rigid structures at this value. **B:** A multiple sequence alignment (MSA) of 7 representative CIN85 homologs shows the exceptional sequence identity between residues 200-239 of the human homolog. The MSA was generated with ClustalOmega ^29^ based on a BLAST ^28^ search of the CIN85 Uniprot entry Q96B97-1 (referenced to residues 200-239). The sequence conservation was plotted as bars on the top of each amino acid position.

It resembled the consensus sequence for PRM recognized by CIN85 SH3 domains (PXXXPR) ^23^ but contained an additional arginine residue (^223^PIKL**R**PR^229^). There is no known PRM in CIN85 N-terminal to this motif, which is why we refer to it as “CIN85-PRM1” in the following.

We further used NMR titrations to assess the interaction of CIN85-PRM1 with isolated SH3 domains of CIN85, observing chemical shift perturbations (CSP) in the ^15^N-labeled SH3 domains upon titration with a synthetic 14-residue peptide of the sequence ^219^FKDKPIKLRPRSIE^232^ (see Supplementary Figure S1). All three domains displayed moderate to weak affinity, with SH3C showing 3 and 5-fold lower dissociation constants (K_D_) than SH3A and SH3B, respectively (K_D_∼0.2-1.1 mM, see Table 1). This suggested the SH3C domain to be the dominant interaction partner to CIN85-PRM1. We mapped the CSP onto the first model of the NMR structure of SH3C (PDB: 2k9g) in Figure 2A. Binding to the CIN85-PRM1 peptide seems to occur through the conserved binding site, involving the RT-loop (residues 278-292), the N-Src-loop (residues 300-307) and also the 3_10_-helix (residues 320-322) of SH3C (see Figure 2B).

**Table 1.**
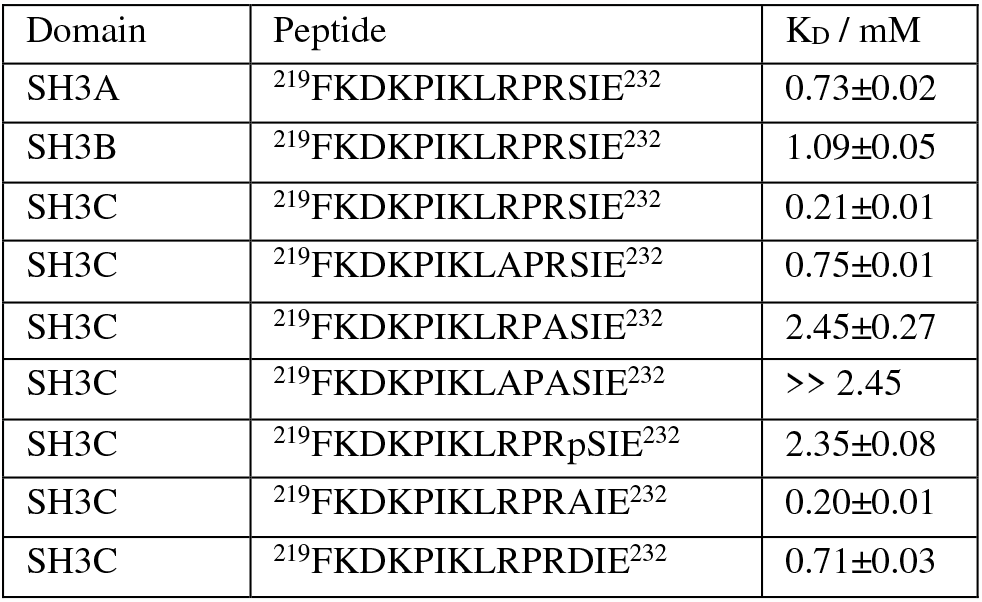
Dissociation constants K_D_ of the binary SH3:peptide interactions as determined by NMR titrations. The error in the fitted K_D_ values was determined via a bootstrap resampling approach.

**Figure 2:**
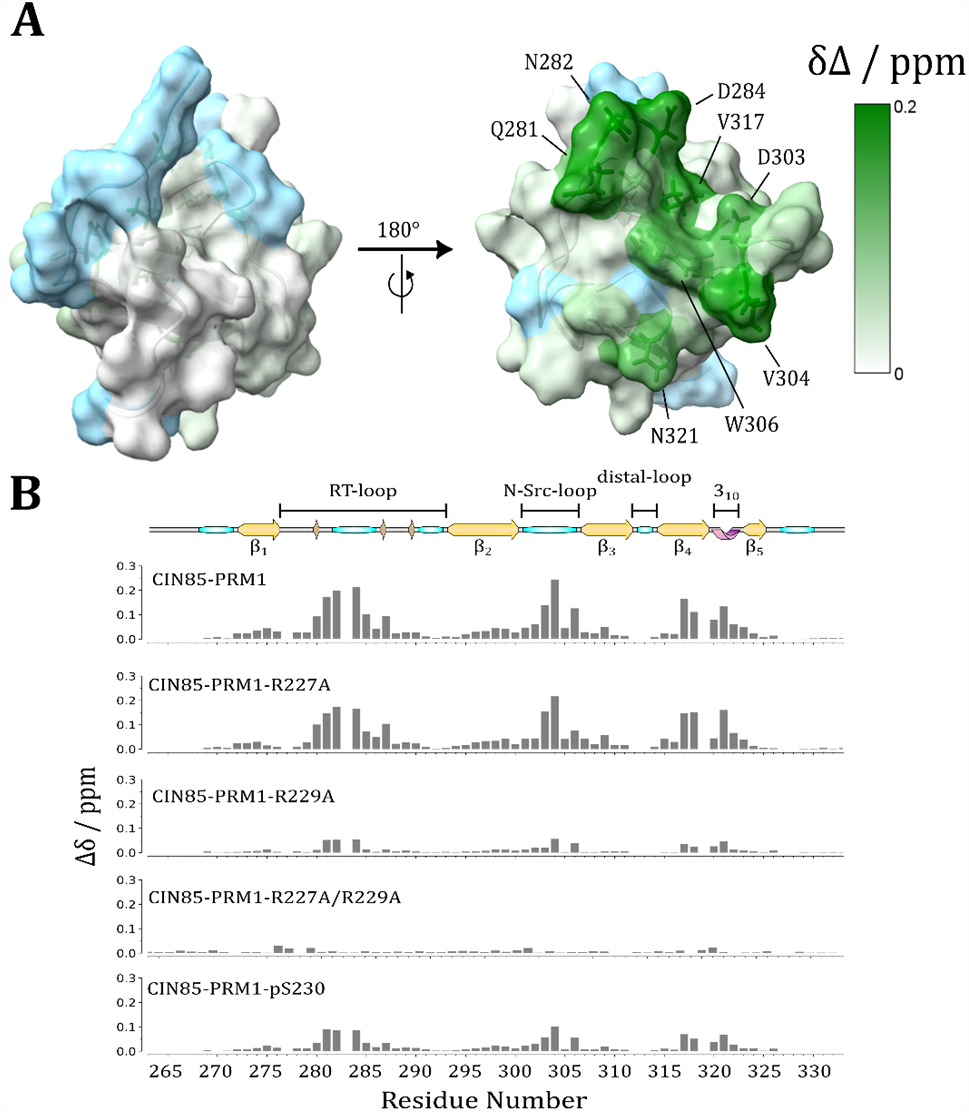
Interaction of CIN85 SH3C to CIN85-PRM1 peptides. **A:** Chemical shift mapping of residues inside the SH3C domain exhibiting CSP when titrated with an excess of the wild-type CIN85-PRM1 peptide. The SH3C domain structure used here was the first structure of the NMR ensemble deposited in the PDB as entry 2k9g. **B:** Bar plots showing the CSP of the residues within the SH3C domain in response to titration with the indicated CIN85-PRM1 peptides. For all the data shown, the molar Ligand:Protein ratio was chosen to be similar, ranging from 10.8-12.9. The secondary structure graph of SH3C on top of this Figure was generated from the STRIDE ^36^ prediction of the PDB entry 2k9g using the SSS-Drawer python script (https://github.com/zharmad/SSS-Drawer).

The recognition of PRM by SH3 domains typically depends on the presence of positively charged residues, such as arginines at the motif’s N-or C-terminus, which often form cation-π interactions with conserved tryptophan residues in the SH3 binding interface ^30^. In past studies on similar systems, introduction of R/A mutations was found to be effective in perturbing this type of interaction ^15, 17, 23, 31^. We therefore used CIN85-PRM1 mutant peptide binding to the SH3C domain to determine the role of both arginines (R227 and R229) in this interaction. The K_D_ increased 4-fold for the R227A and 12-fold for the R229A mutant, with complete abolition of binding only after mutating both residues (see Figure 2B, Table 1 and Supplementary Figure S2). The two arginine residues therefore contribute to the interaction to a different extent, with R229 playing a larger role. Since we knew that this interaction was in part driven by cation-*π* interactions, we anticipated a net energetic contribution on the order of -12 ± 6 kJ/mol if such an interaction would be disturbed by the mutation ^32^. Indeed, we found this difference to be 3.28 ± 0.20 kJ/mol for the R227A and 6.32 ± 1.00 kJ/mol for the R229A mutant peptides. The difference in binding energy compared to wild-type peptide was consistent with loss of a weak cation-π interaction for the R229A mutant, while for R227A, a specific interaction was unlikely, and this residue is more likely involved in non-specific interactions within the binding interface. We suggest that both R227 and R229 tune the interaction in synergy, with R229 contributing most to the binding affinity and thus potentially involved in a more persistent, specific interaction. Interpreting these results, one must bear in mind that these differences in binding energy come from NMR titrations of a synthetic peptide with an isolated SH3 domain. In the context of the whole protein, local concentration and/or co-operative effects could increase the actual strength of this interaction considerably.

The residue S230 is adjacent to R229 and a known site for activation-induced phosphorylation in CIN85 ^33-34^ (see also Supplementary Figure S3). To investigate the effect of S230’s involvement in this interaction, we incorporated a phosphoserine (pS230), a S230D and a S230A mutation into the synthetic peptide. We observed a similar increase in the K_D_ for pS230 as for the R229A mutation, while it was not at all affected by the S230A mutation (see Figure 2B and Table 1). This was consistent with a role of this residue in tuning the extent of interaction to CIN85-PRM1 only by post-translational modification while not being involved in the interaction in general. The phosphomimetic mutation S230D led to a much smaller increase in K_D_, suggesting a specific role of the phosphoryl group in terms of electronegativity and excluded volume ^35^. By correlating the CSP of mutated peptides and the wild-type peptide we also determined that the mode of interaction was conserved, and differences in dissociation constants were only due to weakening of the interaction (Supplementary Figure S4).

### CIN85-PRM1 forms helical structures upon interaction to SH3C

To study the potential interaction of the SH3C domain with CIN85-PRM1 in more detail, we needed to assign the linker backbone resonances first. We accomplished the near-complete assignment of the linker region containing CIN85-PRM1 by acquiring three-dimensional ^13^C-detected experiments on a shorter construct of the CIN85 protein (CIN85_163-333_, see Supplementary Figure S5 for the different protein constructs used in this work). Details of the resonance assignment process, including the assigned ^13^C, ^15^N CON spectrum of CIN85_163−333_, can be found in the Supplement to this article. Within the disordered linker region (residues 163-263), we were successful in assigning 97% the backbone resonances, including the prolines inside CIN85-PRM1. Notably, some of the cross peaks belonging to the core of CIN85-PRM1 were missing from the 3D spectra (L226N-K225C, R227N-L226C and R229N-P228C).

This was likely due to intermediate-exchange line broadening due to the interaction of SH3C with the PRM. This is illustrated here by the signal/noise ratio (SNR) of cross peaks in the ^13^C-detected HNCO spectra of CIN85_163−333_ and of two arginine mutants showing reduced interaction to SH3C in the titration experiments above (R229A and R227A/R229A, see Figure 3C). The resonances within CIN85-PRM1 were severely broadened in both CIN85_163−333_ and the R229A mutant. Only upon introducing the R227A/R229A mutation we observed the SNR to increase to the level of the surrounding linker.

**Figure 3:**
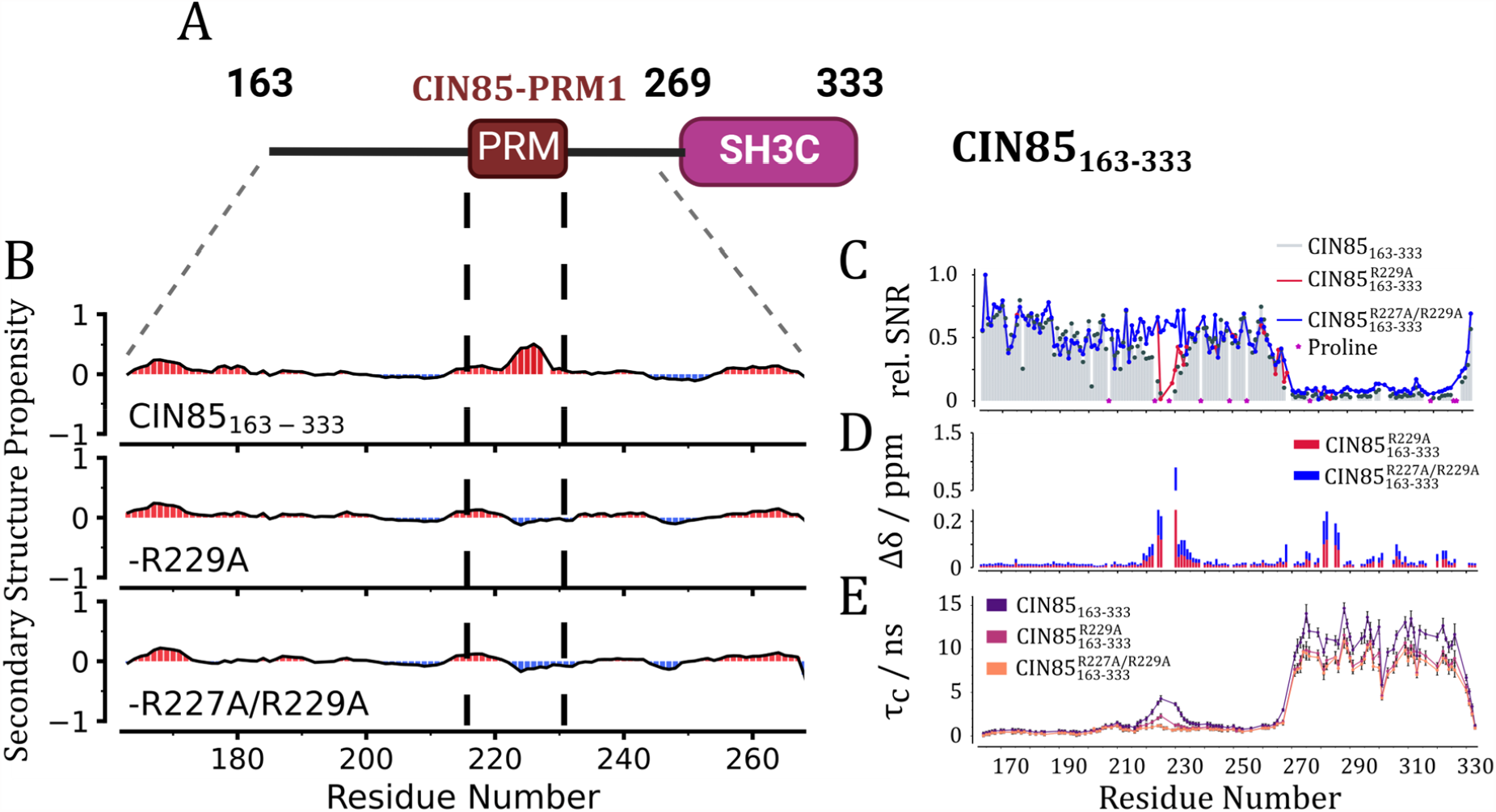
Determination of dynamical and structural properties of CIN85_163-333_. **A:** Domain architecture of CIN85_163-333_. **B:** Secondary structure propensities based on the backbone resonance assignment of CIN85_163−333_ and the two arginine mutants CIN85_163−333_-R229A and CIN85_163−333_-R227A/R229A calculated by using the ncSPC webserver ^37^. Positive values in red indicate propensity for helical structures while negative values report on propensity to form extended structures. **C:** Relative signal/noise ratios of cross-peaks from the ^13^C-detected HNCO spectra normalized to residue I164 for CIN85_163−333_ (grey bars), CIN85_163−333_-R229A (red line) and CIN85_163−333_-R227A/R229A (blue line). Proline residues were marked with magenta stars, since they did not give rise to signals in the ^13^C-detected HNCO spectra. **D:** CSP of cross peaks from the ^13^C-detected HNCO spectra for CIN85_163−333_-R229A (red) and CIN85_163−333_-R227A/R229A (blue), compared to the CIN85_163−333_ chemical shifts. **E:** Residue-specific rotational correlation time of the three CIN85_163−333_ constructs determined using the TRACT experiment ^38^. The protein samples were uniformly ^13^C/^15^N-labeled at a concentration of 1 mM. All experiments were acquired at 800 MHz and a temperature of 298 K.

We used the backbone resonance assignments to compare the chemical shift perturbations (CSP) in the mutants to the wild-type protein. Significant perturbations were observed within CIN85-PRM1 (residues 280-285) and the RT-loop (residues 300-306) of the SH3C domain, consistent with the NMR titration results of CIN85-PRM1 to the SH3C domain shown above (see Figure 3D, 2B). Additionally, the secondary structure propensities (SSP) were predicted using the ncSPC webserver ^37^ and showed a distinct propensity for helical structures within CIN85-PRM1. This was lost upon introduction of the R/A mutations (Figure 3A). The helical structures likely form through a disorder-to-order transition upon binding to SH3C and are consistent with known structures of SH3 domains bound to their respective peptides (Supplementary Figure S7) ^39-40^.

### Small bound-state population of CIN85163-333 based on local correlation times

We further used the TRACT ^38^ experiment to determine residue-specific apparent rotational correlation times (τ_c_) within CIN85_163−333_ using NMR and sample the interaction between SH3C and CIN85-PRM1 (Figure 3E). We observed small local correlation times τ_c_ for linker residues outside of CIN85-PRM1 and an increase within its core for CIN85_163−333_ (τ_c_ = 4.3 ns). The R229A mutant showed a distinct decrease in τ_c_ of residues within CIN85-PRM1 (τ_c_ = 2.3 ns) and for the R227A/R229A mutant there was no significant difference to the surrounding linker (τ_c_ = 0.5-1 ns). Here, we focused on the residues within CIN85-PRM1 with the highest correlation time, as these determine the bound fraction. The smaller correlation times of other residues within the motif can be explained by the individual binding behavior of the amino acids and the variability in complexes formed through fuzzy interactions. Clearly, even in wild-type CIN85_163−333_ none of the residues of CIN85-PRM1 showed the correlation time of the SH3C domain, averaging at around 11 ns. Bound fractions were estimated based on maximum local correlation times in CIN85-PRM1 and the average correlation time of the SH3C domain was calculated to be 31% for CIN85_163−333_, 14% for the R229A mutant, and zero for the R227A/R229A mutant. For an isolated SH3 domain at room temperature, global τ_c_ values are generally found between 4-5 ns ^41^. The elevated τ_c_ for the SH3C domain can be explained mainly by the presence of the linker, as a control experiments with the isolated SH3C domain showed distinctly lower apparent τ_c_ in its absence (Supplementary Figure S8). The interaction between CIN85-PRM1 and the SH3 domain led to another small increase in the apparent correlation time within the SH3C domain compared to the R227A/R229A mutant where binding is abolished (see Figure 3E).

### Arginine sidechain rotational dynamics indicate competition between both arginines in CIN85-PRM1

Since the arginine residues were arguably playing a major role in this interaction, we further probed rotational dynamics of the arginine guanidinium groups in CIN85_163−333_ through multiquantum chemical exchange saturation transfer (MQ-CEST) ^43^ experiments. These allow to sample the restricted rotation of the guanidinium group mediated by noncovalent interactions such as salt bridges and cation-π interactions (Supplementary Figures S9-S11). Unlike R176, R227 and R265 in the disordered linker and R314 and R315 in the SH3C domain, no N^ε^-H^ε^ cross peak was observed for the side-chain of R229, consistent with its involvement in a salt-bridge or cation-π interaction resulting in signal broadening (Supplementary Figure S9). The rate of rotation (k_ex_) around the C^ζ^-N^ε^ bond of the free arginine guanidinium group was 397 ± 4 s^-1^, in close agreement with previous reports ^43-44^. The obtained k_ex_ rates for R176 and R227 were smaller than free arginine but larger than those of R314 and R265/R315, indicating the less restricted rotational dynamics of arginine side-chains in the disordered linker region than in the folded domain (Supplementary Figure S11A). Upon removal of R229 in the CIN85_163−333_-R229A mutant, a small but significant reduction in k_ex_ was observed for R227, while no significant change in k_ex_ was detected for the other arginines. The chemical shift separation Δω between the two N^η^ nuclei exhibited a similar trend, increasing significantly only for R227 in the R229A mutant compared to wild-type CIN85_163−333_ (Supplementary Figure S11B). This intriguing observation may suggest a degree of competition between R227 and R229 in the interaction with SH3C, so that the partial interaction of R227 with SH3C becomes possible only when R229 is absent. The comparatively small effect of the R229A mutation on R227 arginine sidechain rotational dynamics can be reconciled by the low bound fractions for both constructs determined via the TRACT experiments above. As k_ex_ is a population-averaged value, we would expect the difference to increase with the population of the bound state.

### Effective concentration effects favor the interaction of SH3C to CIN85-PRM1

For recognition motifs tethered to their receptor, effective concentration (c_eff_) effects have been shown to have a significant influence on binding ^45-46^. To understand the role of c_eff_ in the SH3C:CIN85-PRM1 interaction, we estimated c_eff_ of SH3C at CIN85-PRM1 using protein:peptide-complex structures predicted by HADDOCK ^47-48^. We used residues with significant CSP in peptide titration experiments to guide the docking process and measured the distance between the last residue of the folded domain and the beginning of the binding motif to obtain the relevant distance in the complex structure based on the approach developed by Kjaergaard *et al*. ^49^. The effective concentration of the predicted complexes was found to be in the range of 1.3-6.3 mM, while the K_D_ obtained from titration of the untethered CIN85-PRM1 peptide to SH3C was 0.2 mM, indicating well-saturated binding in all complexes (Supplementary Figure S12 and S13). The population of the bound, i.e. auto-inhibited, closed state calculated from these effective concentrations was between 0.86 and 0.97. Since the three SH3 domains are highly related, we further assumed the same relevant distances as for SH3C should be applicable to the potential intramolecular complexes between these domains and CIN85-PRM1. We found c_eff_ to range between 0.9-1.3 mM for the SH3A domain and 1.6-4.1 mM for the SH3B domain. This translated to a factor of 30 (SH3C) > 4 (SH3B) > 2 (SH3A) in comparison with the dissociation constants to the free peptide determined above (see Table 1). Therefore, the interaction of SH3C to CIN85-PRM1 is likely to be favored over both remaining SH3 domains based on c_eff_ and K_D._ The small bound-fraction determined above for the wild-type construct (about 31%) was inconsistent with the expected results from the c_eff_ calculations here. However, it is known that disordered regions can behave differently depending on whether they are tethered to folded domains on one or both ends ^50^. Following this, the removal of the SH3B domain from the N-terminal end of CIN85_163−333_ could have increased the entropic chain character of the remaining linker. This would impart a significant increase of conformational freedom on CIN85-PRM1 located in the center of the linker, making the linker as a whole act as an ‘entropic bristle’ and thus making the interaction with SH3C less likely ^45, 51-52^.

### The SH3C domain in CIN851-333 is auto-inhibited by binding to CIN85-PRM1 intramolecularly

To test whether the population of the bound state was indeed higher in CIN85_1-333_, we transferred the assignment of the linker region from the truncated CIN85_163−333_ by comparing their ^1^H−^15^N TROSY spectra (Supplementary Figure S14 and S15) and determined the apparent τ_c_ values for CIN85_1−333_ and its R/A mutants (Figure 4B). The residue S230 was used here as a proxy for the effective *τ*_*C*_ of the CIN85-PRM1 core, since residues L226-R229 could not be assigned in the ^1^H-^15^N TROSY spectrum of CIN85_1-333_. Indeed, we observed a significantly higher population of the bound state in CIN85_1−333_, characterized by a distinct increase in *τ*_*C*_ of residue S230 (τ_c_^S230^ =18.3 ns at 0.5 mM). As for CIN85^163-333^, a R229A or R227A/R229A mutation was effective in abolishing this interaction. In comparison, the median τ_c_ for the three SH3 domains at this concentration ranged between 12.7-18.2 ns (Supplementary Figure S16). Some residues such as D303 (part of the n-Src-loop of SH3C, see also Figure 2B) exhibited values significantly larger than these median values (τ_c_^D303^ = 20.9 ns at 0.5 mM). The elevated τ_c_ observed for S230 can therefore be explained by the interaction of CIN85-PRM1 with the binding interface of one of the SH3 domains.

**Figure 4:**
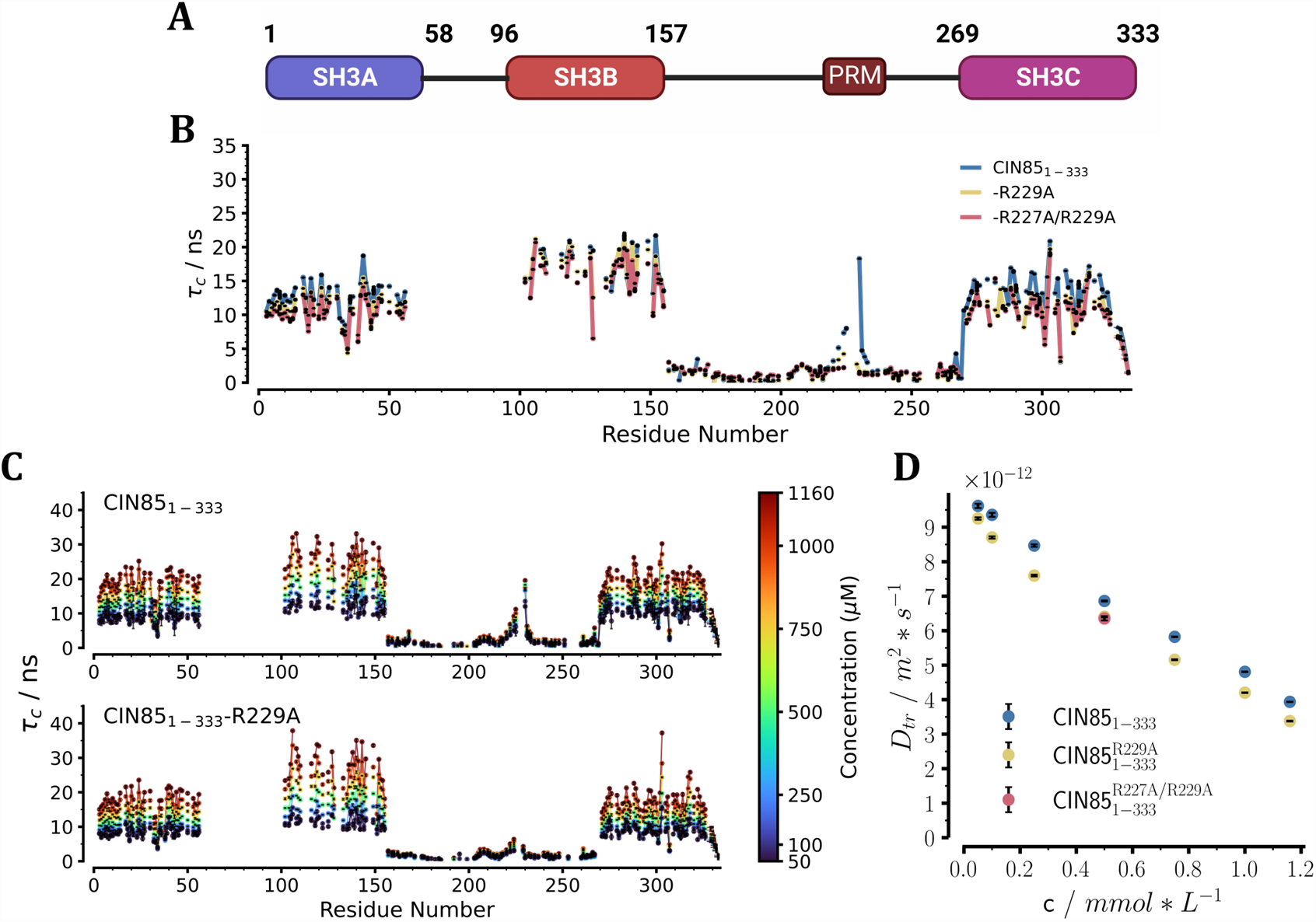
Determination of intermolecular non-specific transient interactions as well as intramolecular specific interactions in CIN85_1-333_. **A:** Domain architecture of CIN85_1-333_.**B:** Residue-specific rotational correlation time of the three CIN85_1−333_ constructs at a molar concentration of 0.5 mM using the TRACT experiment ^38^. **C:** Residue-specific apparent rotational correlation times of CIN85_1−333_ and CIN85_1−333_-R229A in dependence of protein concentration ranging from 0.05 mM to 1.16 mM. **D:** Translational diffusion coefficient D_tr_ of CIN85_1−333_ (blue) and CIN85_1−333_-R229A (yellow) determined using the N-TRO-STE experiment ^42^ in dependence of protein concentration ranging from 0.05 mM to 1.16 mM. The R227A/R229A mutant of CIN85_1-333_ (red) was sampled at a single concentration (0.5 mM). The CIN85_1-333_ constructs used here were expressed uniformly ^15^N labeled and perdeuterated. All experiments were conducted at 800 MHz and 298 K.

To determine whether the interaction with CIN85-PRM1 was dominated by an intramolecular or intermolecular binding mode, we further analyzed residue-specific τ_c_ values and translational diffusion coefficients D_tr_ over a wide concentration range for CIN85_1-333_ and its R229A mutant (Figure 4C+D). The R229A mutant was chosen as a control because it already sufficiently perturbed the interaction with CIN85-PRM1 (Figure 4B). In CIN85_1-333_, we found the ratio of τ_c_ values between residues within CIN85-PRM1 and the SH3 domains independent of concentration, proving that the interaction was intramolecular. As expected, the R229A mutant showed small τ_c_ values within CIN85-PRM1 at all concentrations, more similar to the surrounding linker. These findings were corroborated by the translational diffusion coefficients (D_tr_) determined via N-TRO-STE experiments ^42^ (Figure 4D).

CIN85_1-333_ consistently showed higher D_tr_ values than the R229A mutant did at all concentrations, indicating slower translational motion of the mutant (6%-14% difference, depending on concentration) and consequently a more compact shape of CIN85_1-333_ compared to CIN85_1-333_-R229A.

In addition to the differences between constructs indicating the intramolecular SH3-PRM association, we also observed effects common to both CIN85_1-333_ and the R229A mutant. These showed a general concentration-dependence of local apparent τ_c_ within the SH3 domains and the global *D*_*tr*_ (Figure 4C+D). We also found the slope of the concentration-dependence of *D*_*tr*_ for both constructs to be the same within the experimental uncertainty (Supplementary Figure S17). This showed the concentration-dependence of these parameters to be independent of the interaction to CIN85-PRM1 and thus common to both constructs. To rule out the possibility that viscosity changes with protein concentration were mainly responsible for the observed differences, we measured ^17^O T_1_ relaxation rates of the buffer at different protein concentrations and estimated the resulting viscosity changes ^53^ (Supplementary Figure S17). The measured dynamic viscosity increased by a factor of 1.32 from pure buffer to a CIN85_1-333_ concentration of 1.3 mM. However, the median rotational correlation time of the three SH3 domains in CIN85_1-333_ and the R229A mutant increased by a factor of 1.9-2.3 and 2.0-2.6, respectively (Supplementary Figure S16). Consistently, we found this change to be a factor of 2.4 and 2.7 for D_tr_ (Figure 4D). Therefore, the observed differences in τ_c_ and D_tr_ with concentration cannot be attributed to viscosity changes alone. These differences can only be explained by assuming an increasing extent of transient non-specific protein-protein interactions with concentration ^54-55^, which however do not involve the CIN85-PRM1:SH3 interaction. Thus, in addition to the specific trimerization via the coiled-coil domain that has been described previously ^17^, CIN85 SH3 domains can also mediate non-specific low-affinity oligomerization by themselves.

### CIN85-PRM1 phosphorylated at S230 provides an activation-induced release of CIN85 SH3 domains

The serine residue at position 230 was shown above to significantly weaken the CIN85-PRM1:SH3 association in its phosphorylated state (Figure 2B). In addition, it was found to be highly phosphorylated in a multitude of phosphoproteomic studies (Supplementary Figure S3). In particular, S230 has been shown to be phosphorylated in response to BCR engagement ^33^, which is why we decided to assess the signaling function of this residue. We introduced fluorescently labeled full-length citrine-CIN85 and citrine-CIN85-S230A into DG75 B cells expressing no endogenous CIN85. The intact BCR-related signaling machinery made this cell line suitable for studying the effect of the mutation on B cell signaling ^33^. The S230A mutation was selected because it did not affect SH3C’s interaction with the corresponding mutant peptide, and it created a mutant that could not be phosphorylated at position S230 (see Table 1). It has been shown previously that the multivalent and promiscuous interactions between CIN85 SH3 domains and SLP65 PRMs drive their formation of liquid-like condensates that are a necessary prerequisite for BCR signaling events to occur ^18^. Monitoring the propensity for LLPS is therefore a viable readout for the overall BCR-related signaling capabilities of the S230A mutant. Utilizing imaging flow cytometry, we observed a decrease in the percentage of droplet-positive cells, and attenuated BCR-induced recruitment to the plasma membrane of the S230A variant compared to wild-type CIN85 (see Figure 5A+B). These findings correlated with compromised BCR-induced Ca^2+^ mobilization in CIN85 S230A-expressing DG75 B cells compared to cells expressing the wild-type protein (Figure 5C). Hence, this indicated that phosphorylation at S230 contributes to efficient BCR-induced signaling via liquid-like condensates, corroborating what has been shown previously by Wong *et al*.^18^. We further used label-free mass spectrometry to investigate, how phosphorylation of CIN85-PRM1 changes the CIN85 interaction network. For this purpose, we purified CIN85 from lysates of DG75 B cells expressing either wild-type citrine-CIN85 or citrine-CIN85-S230A, in both the resting and stimulated state (see Supplementary Figure S19 and S20 and Supplementary Table S3). This approach did not reveal a specific protein interacting with CIN85-PRM1. Nevertheless, we observed more subtle differences in the interactome of CIN85. We determined a decreased abundance of both SLP65 and CIN85 in preparations of the S230A variant. Since we did not detect a difference in the expression of the mutant compared to the wild type (Supplementary Figure S21), this can likely be attributed to the reduced propensity for these two proteins to engage in network formation leading to liquid-like condensates (see Figure 5A).

**Figure 5:**
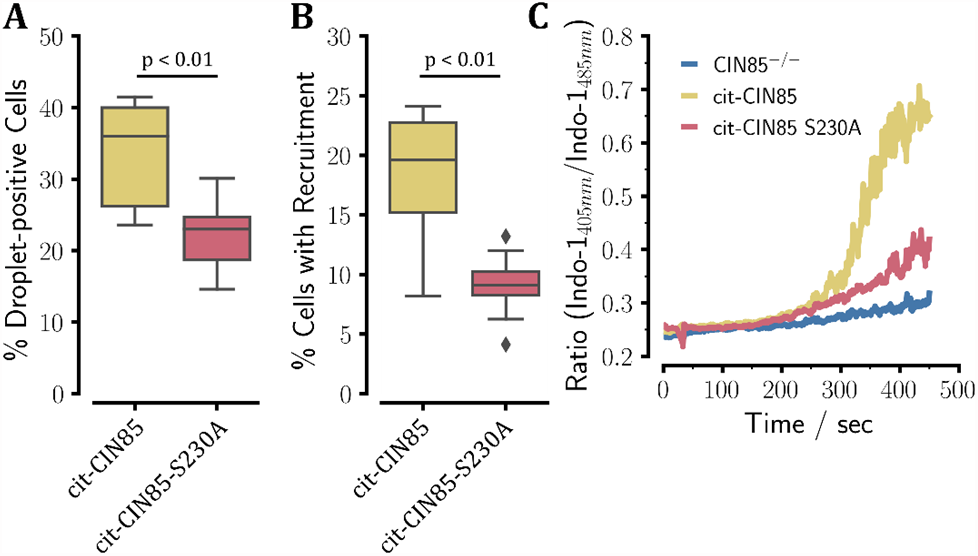
Imaging flow-cytometric characterization of DG75 B cells carrying citrine-CIN85 or citrine-CIN85-S230A. **A:** Percentage of DG75 B cells that showed cytosolic droplets. **B:** Percentage of DG75 B cells that exhibited significant recruitment to the plasma membrane. **C:** Ca^2+^ mobilization sampled as the ratio of the Indo-1 absorbance ratio 405 nm*/*485 nm using DG75 B cells carrying either cit-CIN85(yellow), cit-CIN85-S230A (red) or CIN85^−*/*−^ knockouts (blue).

## Discussion

The interaction networks mediated by scaffold proteins such as CIN85 and SLP65 have been shown to be of primary importance for the physiological signaling processes occurring in human and murine B cells ^56-57^. Previous work addressed the assembly of these two scaffolds into liquid-like pre-signaling clusters that are a prerequisite for the proper function of BCR signaling and enable a rapid cellular response upon the activation of the BCR ^15-18^. Yet, open questions still exist regarding the necessary modifications in CIN85 and SLP65 to modulate their propensity for undergoing LLPS and to enable a signaling-competent state. The CIN85 protein serves a multitude of different context-sensitive functions in a variety of different cell types, which require a finely balanced and tightly controlled regulation. In multi-domain proteins, an effective way of providing regulation by the cell is through a SH3-mediated intramolecular association involving IDRs. This recurring theme of auto-inhibited SH3 domains has been observed for several different systems in immune cells, including the Nck adaptor protein in T cells ^58^ and the cytosolic component of the NADPH oxidase p47^phox^ in phagocytes ^59^. From the perspective of cellular signaling, regulation by intramolecular interactions has the advantage of being concentration independent and, due to effective concentration effects, needing only moderate nominal affinities to compete with potential intermolecular binding partners. Based on our findings, we therefore suggest that one of the SH3 domains is autoinhibited by the intramolecular interaction to CIN85-PRM1 in the adjacent linker region. From the nature of the NMR relaxation experiments presented here (Fig4B and C), we can only definitively say that the bound state is the major populated state, but not whether one or multiple of the domains enforce this interaction synergistically. However, we observed a clear hierarchy of binding affinities of the different SH3 domains to CIN85-PRM1, with SH3C binding the strongest, followed by the SH3A and SH3B (see Table 1). This hierarchy is also supported by the phylogenetic origin of these domains, as SH3B and SH3C have split from the common progenitor SH3 domain first, while the SH3A domain was later generated via gene-duplication of SH3C, making them more similar compared to SH3B ^60^. This also reflects the fact that SH3A and SH3B exhibit strikingly dissimilar binding mechanisms to similar peptides ^61^. In addition, our analysis of dissociation constants and effective concentrations suggests that SH3C is likely to outcompete SH3A and SH3B (see Table 1 and Supplementary Figures S12 and S13). Therefore, based on the data at hand, we argue that it is the SH3C domain that is able to compete with SLP65 PRM’s and is fully occupied by this interaction while the SH3A and SH3B domains are free to engage in these interactions. The given state of a multi-domain scaffold protein like CIN85 is typically determined by the presence or absence of many different nominally low-affinity interactions. This auto-inhibitory interaction between the SH3C domain and CIN85-PRM1 is thus a potential candidate for shifting these equilibria towards a signaling-competent state of the CIN85 protein upon engagement of the BCR. Having established the auto-inhibition of the SH3C domain *in vitro*, we addressed its potential signaling consequences in the cellular context. Since the promiscuous interactions of CIN85 SH3 domains with SLP65 PRM drive the LLPS in conjunction with small vesicles ^18^, modulating the valency of the CIN85 protein should also have an effect on the propensity for phase separation. Indeed, a study conducted in parallel by our group has shown by *in vitro* droplet reconstitution assays and lattice-based computational modeling that the critical concentration for LLPS was lowered for the CIN85_1-333_ mutant in which the interaction was abolished ^31^. This can be only understood quantitatively if one assumes the presence of an autoinhibitory interaction, preventing one of the SH3 domains to interact with SLP65 PRMs. These results are therefore consistent with the proposed autoinhibition of the SH3C domain, as discussed above. The activation-induced phosphorylation of S230, located directly adjacent to CIN85-PRM1 (Figure 1B), has been shown to occur shortly after the stimulation of the BCR ^33^. Taking into account the presented evidence on this intramolecular interaction *in vitro*, we suggest that this motif acts as a “switch” to modulate CIN85’s valency depending on the cellular signaling state. The data from cultured B cells presented here supports this hypothesis, as the propensity for LLPS was significantly decreased in the citrine-CIN85-S230A mutant cells, where this interaction is expected to be persistently active (Figure 5A). This also suggests a significant population of phosphorylated CIN85-PRM1 in the state of tonic signaling ^62-63^, since this difference was already apparent in the non-stimulated cells. Modulating the extent of this interaction could therefore help to maintain the correct level of LLPS in resting B cells and shift the equilibrium, depending on their cellular activation state. This in turn can influence the response of these cells to external stimuli, which was evident here from the reduced recruitment of CIN85 to the plasma membrane and the diminished mobilization of Ca^2+^ in response to BCR engagement (see Figure 5B and C).

In conclusion, we propose a mechanism by which the SH3C domain of CIN85 is autoinhibited by an intramolecular interaction to CIN85-PRM1. This interaction is regulated intracellularly through phosphorylation at the neighboring residue S230, which upon modification enables the SH3C domain to engage in interactions to SLP65 and other effectors, promoting the physiological signaling-competent state. We further propose that the modulation of the interaction between CIN85 SH3 domains and CIN85-PRM1 is important for maintaining the preformed signaling clusters of CIN85 and SLP65 to allow for a dynamic and contextual response, depending on the cellular activation state.

## Associated Content

Experimental section; resonance assignment of flexible linker regions in CIN85 using a combination of ^1^H and ^13^C-detected NMR experiments; determination of intramolecular effective concentrations from HADDOCK-derived complexes; determination of arginine sidechain dynamics using MQ-CEST experiments; Mass-spectrometric analysis of protein abundances in DG75 B cells. Assigned chemical shifts for CIN85_163-333_ constructs.

## Acknowledgements

The authors thank Claudia Schwiegk for the expression and purification of the recombinant protein samples used throughout this work. The authors also thank Kerstin Overkamp for the synthesis of the peptides used in the titration experiments. We are grateful to Gogulan Karunanithy and D. Flemming Hansen for providing the MQ-CEST pulse programs and analysis scripts, and to Joachim Maier for useful discussions.

This paper was typeset with the bioRxiv word template by @Chrelli: www.github.com/chrelli/bioRxiv-word-template

## Author contributions

The manuscript was written through contributions of all authors. / All authors have given approval to the final version of the manuscript

