## Supplementary Information for "Autoinhibition in the Signal Transducer CIN85 Modulates B Cell Activation"

**to the article**

**by Sieme *et al.***

### Supplementary Methods

#### Peptide Synthesis

The peptides used during the course of this work were synthesized by means of solid-phase peptide synthesis with an acetylated N-terminus and amidated C-terminus. A LiChrosorb RP-18 column (MerckMillipore) was used for the purification and peptides were dissolved in demineralized $H_2O$ with 0.1% trifluoroacetic acid (TFA) as a running buffer prior to injection. Elution was performed with a linear gradient of 0-100% (v/v) acetonitrile with 0.1% TFA. The purity was checked to be larger than 95% by liquid chromatography-electrospray mass spectrometry. After lyophilization the peptides were stored at -20°C until needed.

#### Sample preparation for in vitro experiments

$CIN85_{1-333}$, $CIN85_{\Delta 57}$ and $SLP65_{1-330}$ were expressed and purified as described previously [1]. The $CIN85_{163-333}$ construct was generated based on the $CIN85_{1-333}$ construct using the primers 5'-GCG TCT AGA GGA TCC GGC ATT TCC CAG GAT GAG CAG-3' and 5'-GTC CTC GAG AAG CTT TAT TCC TTT TCA AAG TCC GGT GGA AGT AAC-3'. The R227A, R229A, S230A and S230D mutations were then introduced into the corresponding construct by PCR-based site-directed mutagenesis (Quikchange II, Agilent) and expressed and purified with the same protocol as for the first three constructs. The individual SH3 domains were expressed in *Escherichia coli* BL21(DE3) (New England Biolabs). For CIN85 SH3A and SH3B, the coding sequences were cloned into a pGEX4-T vector, which also encoded a N-terminal glutathione-S-transferase (GST) affinity tag and a Thrombin cleavage site. After cell lysis the proteins were purified by affinity chromatography using Pierce®Glutathione Agarose. The final purification step was accomplished by means of size exclusion chromatography using a HiLoad®16/600 Superdex®75 pg (GE Healthcare) chromatography column. The purified protein was then transferred to the final buffer containing 20mM HEPES pH 7.2, 100mM NaCl, 1.0mM TCEP and 0.5mM Pefabloc using dialysis. Aliquots were stored at -80°C. For CIN85-SH3C, the coding sequence was cloned into a modified pET16b vector encoding a N-terminal 7xHis affinity tag and a tobacco mosaic virus (TEV) protease cleavage site. Affinity purification was carried out using a 5mL Ni-NTA Protino®column (Macherey-Nagel) and the final purification step was carried out via size-exclusion chromatography using the same type of column as for SH3A and SH3B. The sample was dialyzed into the same buffer as for SH3A and SH3B and aliquots were stored at -80°C. For samples that required isotopic labeling, bacteria were cultured in M9 minimal medium with $^{15}N$-$NH_4Cl$ and/or $^{13}C_6$-D-glucose as the nitrogen and carbon source, respectively. For samples requiring additional perdeuteration, the expression was carried out in $D_2O$-based M9 minimal medium. For producing fluorescently labeled $SLP65_{1-330}$-S236C/C271A, residue C236 was introduce by PCR-based site-directed mutagenesis (Quikchange II, Agilent). The mutant protein was produced as described above and ATTO 430LS dye (Atto-Tec) was conjugated to C236 by a maleimide thiol reaction (according to the manufacturer´s protocol). The reaction product was purified by size exclusion chromatography (Superdex 75 10/300 GL, GE Healthcare) before usage.

#### Cells, expression vectors, Ca2+ monitoring, and immunopurification

DG75 B cells were cultured in RPMI 1640 supplemented with 10% fetal calf serum, 2 mM L-glutamine, 1 mM sodium pyruvate, 50 µM 2-mercaptoethanol and antibiotics. B cell receptor (BCR) stimulation of the cells was performed with 0.5µg/mL - 2µg/mL of F(ab')2-anti-human IgM (Jackson ImmunoResearch) at 37°C. For the generation of Cbl-interacting protein of 85 kDa (CIN85) and CD2AP-double-deficient cells we used CRISPR/Cas9-mediated genome editing using the sg-RNAs GAT GAA TTA ACT ATT CGA GTT GG (targeting exon 2 of CD2AP) and CCC TGC ACG AAG TGC CCA GTG GA (targeting exon 3 of SH3KBP1). For reconstitution, the cDNAs encoding human CIN85 and variants, respectively, were cloned into the pMSCV vector (Clontech) containing the sequence for the EGFP variant citrine, resulting in N-terminal citrine-tagged proteins, and allowing for retroviral transduction. For flow cytometric monitoring of the BCR-induced $Ca^{2+}$ mobilization, cells were stained with Indo-1 and analysed as described previously [2-3]. For purification of citrine-CIN85 or variants for mass spectrometry analysis, $2 \times 10^8$ cells were lysed in lysis buffer (50 mM TRIS/HCl pH 7.5, 150 mM NaCl, 1.0 mM EDTA, 0.5% Deoxycholate, 0,1% SDS; 1% (v/v) NP40; 1.0 mM $Na_3VO_4$, 1 tablet cOmplete®-EDTA free (Roche)) for 1 h. Samples were cleared by 10 min centrifugation at 20,000 × g. Supernatants were incubated with GFP selector beads (Nano-Tag). Beads were washed 6 times with lysis buffer and treated with lithium dodecyl sulfate (LDS) sample buffer at 70°C for 10 min.

#### Imaging Flow Cytometry

CIN85/CD2AP-deficient DG75 cells expressing citrine-CIN85 and the citrine-CIN85-S230A variant, respectively, were analysed by imaging flow cytometry (Luminex ImageStream X MKII). For evaluating the differences in droplet formation, 3000 cells were monitored in $n > 10$ experiments. For analysis via the Ideas software (Luminex), a mask was generated using the ”features spot” (M=2,Ch2, bright 39 82 3.1) and based on this mask the drop count feature was utilized. The proportion of cells having $n > 0$ of the defined spots was determined. For the analysis of BCR-induced recruitment of CIN85 proteins, 1500 cells were measured without or 3min after BCR engagement. Based on the “feature finder” tool of the Ideas software, recruited cells were identified using the ”spot count (Ch02)” and “H variance mean (Ch02)” features. For further analysis, the proportion of cells with recruited CIN85 was determined based on the “spot count” high and “H variance mean” for $n > 15$ experiments.

#### Mass Spectrometric Analyses

Mass spectrometric analyses were performed by the Core Facility Proteomics at the University Medical Center Göttingen. Samples were reconstituted in 1×NuPAGE LDS Sample Buffer (Invitrogen) and run into 4-12% NuPAGE Novex Bis-Tris Minigels (Invitrogen) for 1 cm distance. Gels were stained with Coomassie Blue for visualization purposes, and each sample cut out as a whole and diced. After washing, gel slices were reduced with dithiothreitol (DTT), alkylated with 2-iodoacetamide and digested with the endopeptidase Trypsin (sequencing grade, Promega) overnight. The resulting peptide mixtures were then extracted, dried in a SpeedVac, reconstituted in 2% acetonitrile/0.1% formic acid/ (v:v) and prepared for nanoLC-MS/MS as described previously [4].

For mass spectrometric analysis samples were enriched on a self-packed reversed phase C18 precolumn (0.15 mm ID x 20 mm, Reprosil-Pur120 C18-AQ 5 µm, Dr. Maisch, Ammerbuch-Entringen, Germany) and separated on an analytical reversed phase-C18 column (0.075 mm ID × 200 mm, Reprosil-Pur 120 C18-AQ, 3 µm, Dr. Maisch) using a 30 min linear gradient of 5-35% acetonitrile/0.1% formic acid (v:v) at 300 nL $min^{-1}$. The eluent was analyzed on a Q Exactive hybrid quadrupole/orbitrap mass spectrometer (ThermoFisher Scientific, Dreieich, Germany) equipped with a FlexIon nanoSpray source and operated under Excalibur 2.5 software using data-independent acquisition (DIA). Each experimental cycle contained one full mass spectrometry

(MS) scan across the 350-1600 *m/z* at a resolution setting of 70000 full width at half maximum (FWHM), and AGC target of $1 \times 10^6$ and a maximum fill time of 60msec, and 11 variable window size MS/MS segments recorded at a normalized collision energy setting of 25%, a resolution setting of 17500 FWHM, an AGC target of $1 \times 10^6$ and a maximum fill time of 60msec. Three technical replicates per sample were acquired.

DIA-MS data were processed using Spectronaut version 16.0 (Biognosys, Schlieren, Switzerland). Proteins were identified using the Pulsar search engine at default settings against the UniProtKB human reference proteome v2021.01 along with a set of common lab contaminants. Protein quantitation was achieved by integrating up to 6 fragments per precursor, and up to 10 precursors per protein at 1.0% false discovery rate (FDR), respectively. Protein abundance values were quartile normalized and used for further statistical analysis.

#### NMR Spectroscopy

##### NMR Titration experiments

For the NMR titration experiments, the isolated SH3 domains were diluted from the concentrated aliquots to the desired concentration into buffer containing 20mmol HEPES, 100mM NaCl, 1mM TCEP, 0.5mM Pefabloc®SC, 5% $D_2O$ and 0.5mM DSS as the chemical shift standard. The samples were finally transferred to 3mm NMR tubes (Hilgenberg GmbH, Malsfeld). The lyophilized peptides were dissolved in 20mmol HEPES, 100mM NaCl and the pH was again adjusted to 7.2, if needed. Standard $^{1}H-^{15}N$ HSQC experiments (Bruker pulse sequence hsqcetf3gpsi2) were collected at every titration point and the peptide was added successively to the sample, ensuring sufficient mixing before acquiring each experiment. 2D spectra were in general apodized by a cosine-squared function in both dimensions and processed using the NMRPipe software [5] (10.9 Rev 2021.258.11.26 64-bit). Dissociation constants were obtained by a global nonlinear least-squares fit of the obtained CSP of residues exhibiting fast-exchange to a two-state binding model with the following equation [6]:

$$\Delta\delta_{obs} = \Delta\delta_{max} \frac{(([P]+[L]+K_D)-([P+L]+K_D)^2-4[P][L])^{\frac{1}{2}}}{2[P]} \quad (1)$$

These calculations were performed in Python 3.7.1 using the lmfit package v1.0.2 [7]. Errors in the resulting dissociation constants were determined by performing a bootstrapping error calculation with 1000 resampled datasets with replacement. After obtaining the dissociation constant, the Gibbs free energy of this interaction was calculated using the following relation:

$$\Delta G^0 = RTln(K_D) \tag{2}$$

with $R$ as the ideal gas constant (8.314 $Jmol^{-1}K^{-1}$) and $T$ as the absolute temperature in K. This quantity is useful when comparing two similar interactions that differ in their $K_D$, i.e. variants of the same peptide that have a single-point mutation or a post-translational modification (PTM) and exhibit differential binding. Their difference in free energy can then be calculated as follows:

$$\Delta\Delta G = G_1^0 - G_2^0 \tag{3}$$

$$\Delta\Delta G = RTln\left(\frac{K_D^1}{K_D^2}\right) \tag{4}$$

The value of $\Delta\Delta G$ here has the meaning of the stability contribution or net energetic contribution of the residue that was mutated.

**Backbone resonance assignment of CIN85 linker regions**

3D experiments for the resonance assignment of CIN85 linker regions were acquired using magnetic field strengths from 700-1200 MHz and spectra were sparsely sampled to 5-10% of the total points using poisson-gap schedules generated via http://gwagner.med.harvard.edu/intranet/hmsIST/gensched_new.html. Spectral reconstruction was carried out using SMILE [8] or hmsIST [9]. Inspection of the processed spectra was done using POKY [10]. The iterative semi-automated assignment was accomplished using FLYA [11] using the custom experiment definitions for the $^{13}C$-detected experiments listed below. The buffer condition for the samples used for the backbone resonance assignment was as follows: 20 mM MES pH 6.0, 100 mM NaCl, 1.0 mM TCEP, 0.5 mM DSS, 5 vol% $D_2O$, 0.5 mM EDTA and 0.01% $NaN_3$.

**Determination of rotational correlation times and translational diffusion coefficients**

Residue-specific cross-correlated relaxation rates were determined at 800 MHz via the 2D TRACT experiment [1, 12-13], using variable delays from 1-190 ms. The number of scans was varied depending on the protein construct and concentration to obtain an acceptable SNR (NS between 8-64). Rates were fitted using a single exponential model. Calculation of apparent rotational correlation time ($\tau_c$) values from the cross-correlated $\eta_{xy}$ rates was done using a published python script from Robson *et al.* [14].

The N-TRO-STE diffusion experiments [15] were acquired at 800 MHz. Translational diffusion coefficients ($D_{\mathrm{tr}}$) were obtained through fitting the pulsed-field gradient-based intensity attenuation data to Stejskal-Tanner (ST) equation using a non-linear least squares approach [15]:

$$I = I_0 \exp[-D_{tr}\mathrm{Q}] \quad (5)$$

with $\mathrm{Q} = \gamma_H^2 g^2 \delta^2(\Delta - \frac{\delta}{3})$. $\gamma_H$ is the gyromagnetic ratio of $^1$H, $g$ is the peak amplitude of the gradient pulse, and $\Delta$ and $\delta$ are the durations of the long diffusion delay and the short bipolar gradient pulses, respectively. $\Delta$ and $\delta$ were chosen as 70 ms and 3.7-5.95 ms, respectively. The buffer condition for the samples used for relaxation and diffusion experiments was as follows: 20 mM HEPES pH 7.2, 100 mM NaCl, 1.0 mM TCEP, 0.5 mM DSS, 5 vol% $D_2O$, 0.5 mM EDTA and 0.01% $NaN_3$.

**Arginine sidechain MQ-CEST experiments**

The arginine sidechain MQ-CEST experiments introduced by Karunanithy *et al.* [16] were acquired for a reference sample of 100 mM free $^{13}C_6$,$^{15}N_4$]-L-arginine (25 mM HEPES, pH 5.1, 10% $D_2O$) and for $\mathrm{CIN85}_{163\text{-}333}$ and its R229A mutant. The buffer condition for the protein samples used for MQ-CEST experiments was as follows: 20 mM MES pH 6.0, 100 mM NaCl, 1.0 mM TCEP, 0.5 mM DSS, 5 vol% $D_2O$, 0.5 mM EDTA and 0.01% $NaN_3$. All experiments were acquired at a magnetic field strength of 800 MHz and at 278 K, using the $^1$H-detected version of the pulse sequence yielding the $^1H^{\varepsilon}$-$^{15}N^{\varepsilon}$ correlation [16]. The $^{15}$N CEST elements ($T_{\mathrm{CEST}}$) were 250 ms long and $^{15}$N $B_1$ fields of 11.8 Hz, 14.6 Hz, 17.33 Hz, 20.1 Hz and 22.9 Hz were used for the free arginine sample. For the protein samples, $^{15}$N $B_1$ fields of 11.8 Hz, 17.3 Hz and 22.9 Hz were used. The $^{15}$N $B_1$ fields were calibrated as described elsewhere [17].

The $^{15}N$ carrier offsets for both the free arginine and the protein samples were chosen to span the chemical shift range between 63.3-80.1 ppm using 69 evenly spaced steps of 20 Hz. One reference spectrum per $^{15}N$ $B_1$ field value was acquired, setting $T_{CEST}$ equal to zero. For the following analysis, peak intensities were quantified using NMRPipe (10.9 Rev 2021.258.11.26 64-bit) [5] and intensity ratios ($I/I_0$) were calculated based on the respective peak intensity in the reference spectrum. The least-square fitting procedure was done according to Karunanithy *et al.* [16], providing rates of rotation ($k_{ex}$) around the $N^{\varepsilon}$-$C^{\zeta}$ bond and the chemical shift difference ($\Delta\omega$) between the two $N^{\eta}$ nuclei of arginine sidechain guanidinium groups. The arginine sidechain $^{1}H^{\varepsilon}$-$^{15}N^{\varepsilon}$ cross peaks were assigned in $CIN85_{163\text{-}333}$ based on the correlations from the backbone amide resonances to the sidechain resonances in a 3D HSQC-NOESY spectrum. The spectrum was acquired using the *noesyhsqcf3gpwg3d* pulse sequence from the standard Bruker pulse sequence library ($\tau_{mix}$ = 150 ms). The size of time-domain was chosen as 2048/64/128 ($^{1}H/^{15}N/^{1}H$) points for the direct and the two indirect dimensions, respectively. The experiment was acquired at 900 MHz and 283 K. NUS was used to reduce the amount of time-domain points that needed to be sampled, amounting to 41% of the full matrix. Spectral processing and NUS reconstruction was accomplished as described above. Analysis of the spectrum was done using POKY [10].

**Resonance Assignment of the IDR containing CIN85-PRM1**

To study the intramolecular CIN85-PRM1/SH3C interaction, observing the binding induced chemical shift changes and expected line broadening of CIN85-PRM1 resonances, their assignments are necessary. We initially used $^{1}H$-detected experiments to assign the complete flexible linker regions in uniformly [$^{13}C/^{15}N$]-labeled $CIN85_{1\text{–}333}$. The different multi-domain constructs of the CIN85 protein used throughout this work are schematically shown in Supplementary Figure S5. Due to extensive line-broadening in the vicinity of CIN85-PRM1 in particular and the high proline content in the linker regions in general, we were not able to assign them completely (25% and 87% sequence coverage for the first and second linker region, respectively). The line-broadening observed specifically in the vicinity of CIN85-PRM1 may be due to exchange with water and between bound and unbound states of the motif. In addition, the bound state is likely experiencing a slowing-down of rotational tumbling, leading to increased transverse relaxation rates. Indeed, the correlations to residues within the folded domains were also not accessible by these experiments due to their large rotational correlation time $\tau_c$.

We addressed this problem in two ways: firstly, by reducing the system size and looking at the linker region of interest separately, and secondly by switching to $^{13}$C-detected experiments that can directly detect proline residues and avoid intermediate-exchange line-broadening effects that are especially strong for $^{1}$H resonances. Additionally, $^{13}$C-detection allowed increasing the temperature from 283 K to 298 K without suffering from increased solvent-exchange.

The construct containing the second linker region, CIN85$_{163-333}$, was stable and allowed for near-complete assignment of backbone and sidechain resonances using 3D $^{13}$C-detected experiments. Despite lacking the SH3B domain at its N-terminus, the majority of resonances were unperturbed compared to the longer CIN85$_{1-333}$ construct (Supplementary Figure S14 and S15). By this approach, 97% of the linker backbone resonances, including most aliphatic sidechains, were assigned. However, this excluded the correlations to the residues L226, R227 and R229 in the 3D spectra, which belong to the core of CIN85-PRM1 and were likely broadened beyond detection (see Figure 3C in the manuscript).

The R229A and R227A/R229A mutations likely perturb the interaction in multi-domain CIN85 constructs, based on their effects observed in SH3C-peptide titrations (Figure 2 in the manuscript). Therefore, we applied the $^{13}$C-detected approach discussed above to assign the CIN85$_{163-333}$ constructs carrying these mutations. The full backbone assignment was obtained for the R227A/R229A mutant, while the $^{1}$H$^{N}$ resonances of L226 and R227 were not accessible in the R229A mutant. The resonance assignment for all three constructs can be found in the supplementary to this article.

The experimental parameters of the triple-resonance experiments utilized in this study can be found in Supplementary Table S1 and Supplementary Table S2.

#### Determination of Effective Concentrations

We utilized the '$c_{\mathrm{eff}}$ Calculator' webserver [18] to estimate this parameter for the SH3C:PRM interaction. We used the backbone and sidechain chemical shifts of the region between residues 163-269 determined for the CIN85$_{163-333}$ construct to generate dihedral angles using TALOS-N [19]. These dihedral angles were then used to generate an ensemble of 30 structures of a peptide ranging from residue 219-232, utilizing the error-distribution in the predicted $\phi/\psi$ angles from TALOS-N to model the heterogeneity of the resulting peptide structure. We then used

HADDOCK [20-21] to dock the peptide to the SH3C domain (pdb: 2k9g), using the residues of the SH3C domain exhibiting significant CSP in the NMR titration experiment as 'active' residues (residues 281, 282, 284, 285, 303, 304, 306, 317, 321, 322). In the peptide, we used both proline and arginine residues as 'active' residues (P223, R227, P228, R229). For the resulting structures in the four best clusters determined by HADDOCK based on the Z-score, distances between residues 232 at the C-terminal end of the docked peptide and the end of the linker at the SH3C domain (K269) were determined. This was based on the approach detailed in Kjaergaard *et al.* [18] for the determination of effective concentrations in an autoinhibited complex. In total, 16 complex structures were analyzed in this way and this resulted in the spread of potential effective concentrations ranging from 1.3-6.3 mM, depending on the corresponding contact distances shown in Supplementary Figure S12. From the derived effective concentrations and the dissociation constant of the interaction between the untethered CIN85-PRM1 and the SH3C domain obtained from the NMR titration experiments (see Table 1 in the manuscript), we calculated the equilibrium constant of this intramolecular interaction $K_{\mathrm{intra}}$ using Equation 6. Following this we further determined the populations of the bound and the unbound state by applying Equations 7 and 8, respectively. The corresponding bar plot illustrating the populations of the two states is shown in Supplementary Figure S11. We found the population of the closed state ranging from 0.97-0.86 going from the highest determined local concentration to the lowest.

$$K_{intra} = \frac{c_{eff}}{K_D} \quad (6)$$

$$f_{bound} = \frac{K_{intra}}{1+K_{intra}} \quad (7)$$

$$f_{unbound} = \frac{1}{1+K_{intra}} \quad (8)$$

### Supplementary Figures

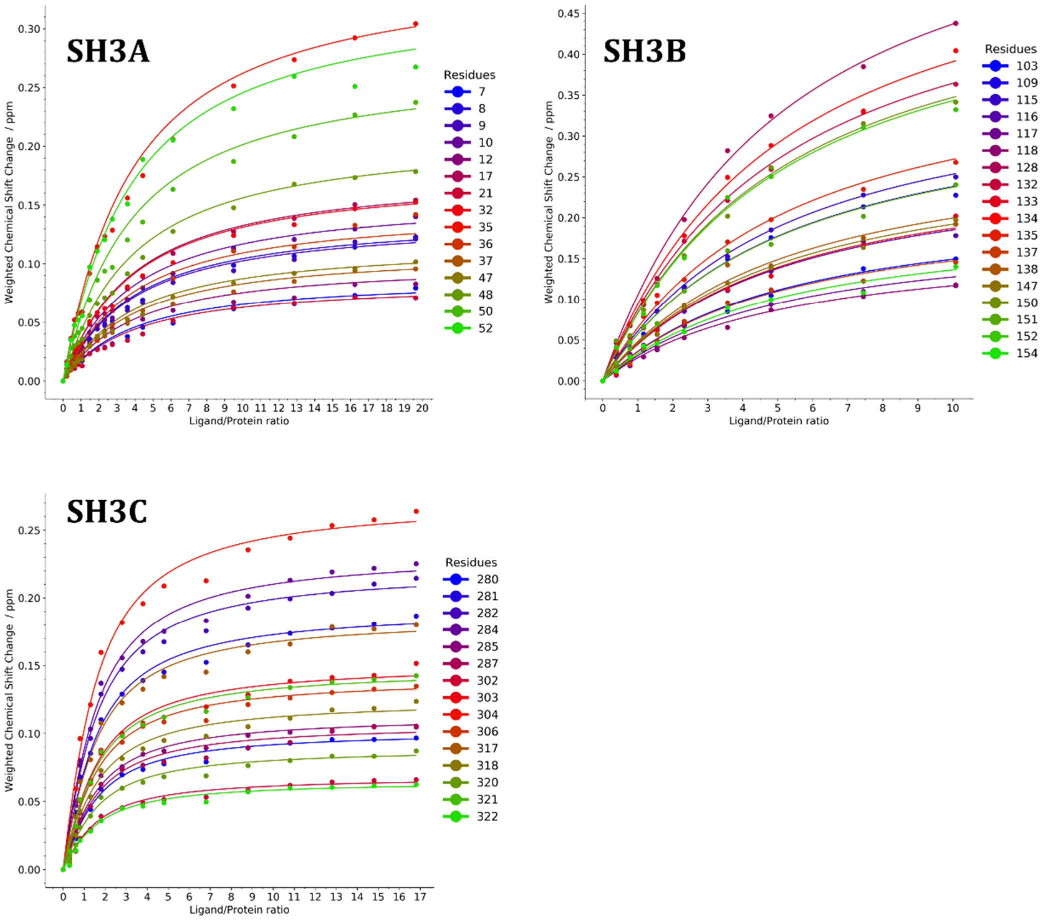


**Supplementary Figure S1:** Global fits of chemical shift perturbation (CSP) curves obtained after titration of $^{15}$N-labeled CIN85 SH3A-C domains with increasing amounts of CIN85-PRM1.

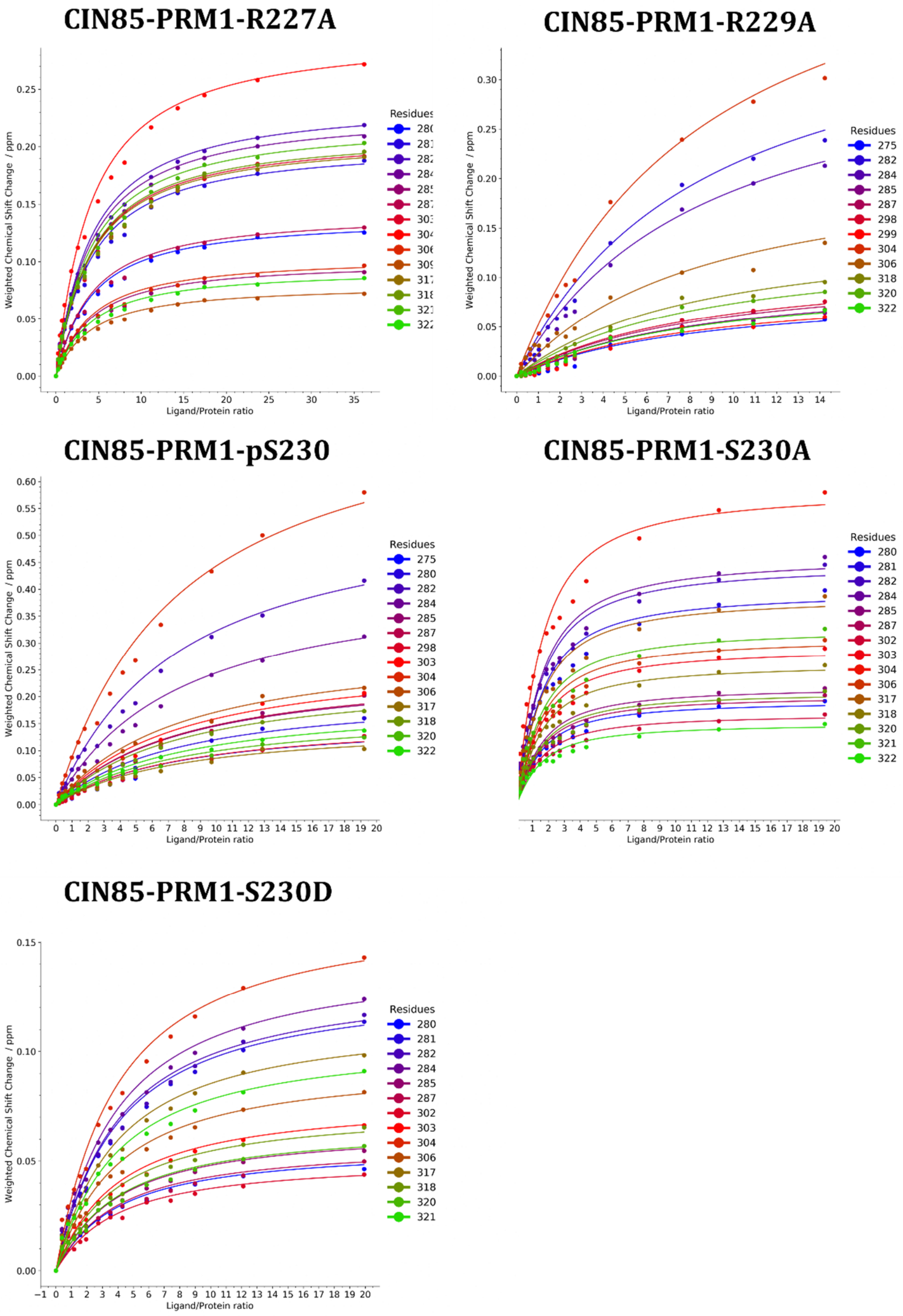


**Supplementary Figure S2:** Global fits of chemical shift perturbation (CSP) curves obtained after titration of $^{15}$N-labeled CIN85 SH3C domain with increasing amounts of CIN85-PRM1 peptides carrying the modifications indicated in the respective insets.

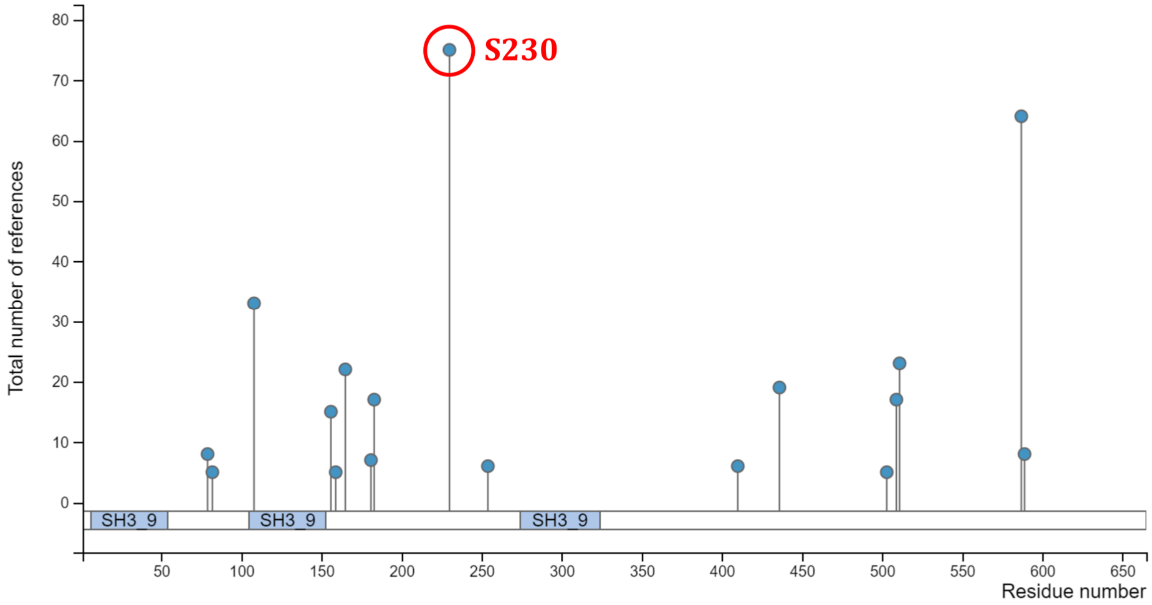


**Supplementary Figure S3:** Lollipop plot showing known phosphorylation sites of human CIN85 as generated by PhosphoSitePlus® v6.6.0.4[83] for SH3KBP1(human) (Accessed on 2022/06/21 at 17:16 CET). The vertical axis shows the number of publications describing the phosphorylation at the corresponding residue number. Only sites are shown that were mentioned in at least 5 independent studies.

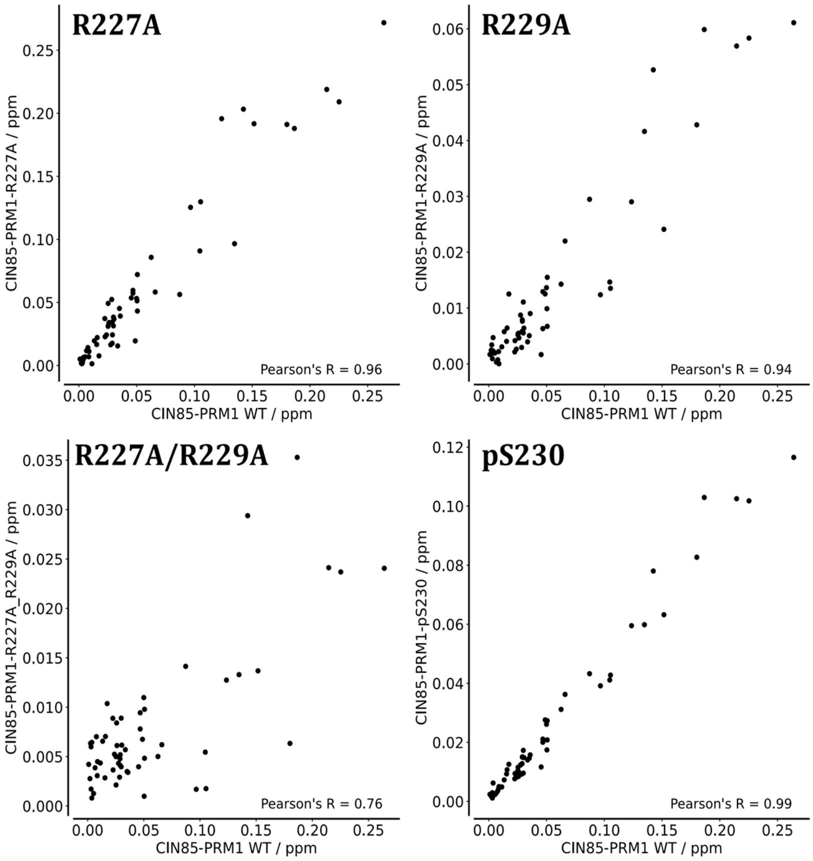


**Supplementary Figure S4:** Correlation plots of CSP observed for residues inside the SH3C domain for the different peptides compared to the one obtained for the titration of this domain with the wild-type CIN85-PRM1 peptide. The correlation was quantified using Pearson's correlation coefficient.

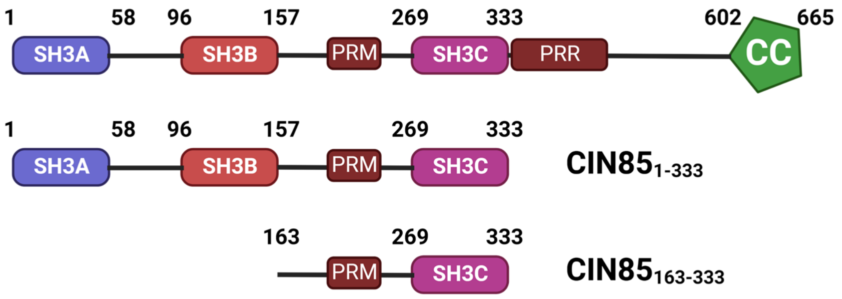


**Supplementary Figure S5:** Domain boundaries of the CIN85 constructs used throughout the study. The numbers indicate the sequence boundaries of the different functional elements of CIN85. PRM and PRR stand for proline-rich motif and proline-rich region, respectively. CC stands for the C-terminal coiled-coil domain. Created via BioRender.com

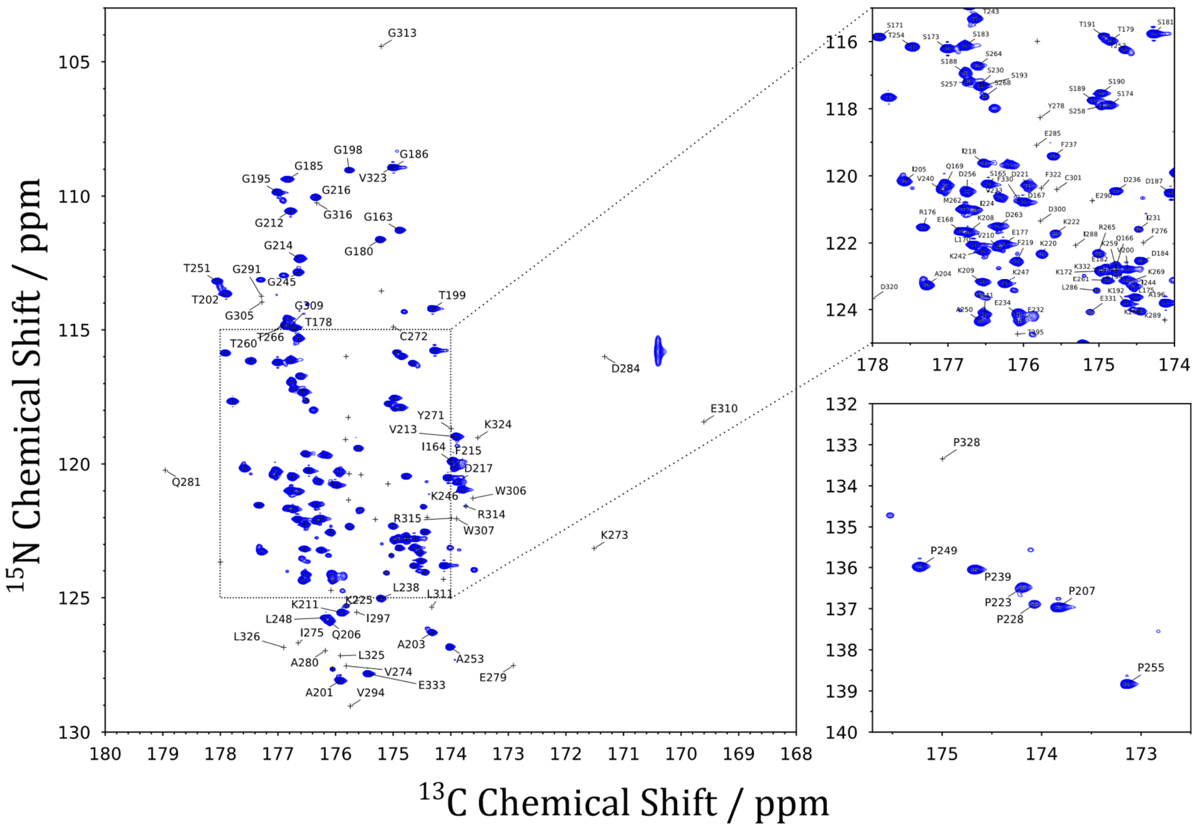


**Supplementary Figure S6: A:** $^{13}C$-$^{15}N$-(HACA)CON spectrum of uniformly $^{13}C$-$^{15}N$-labeled CIN85$_{163–333}$ at 1 mM protein concentration (800 MHz, 298 K) with the annotated backbone resonance assignment. **B:** Zoom-in on the central crowded region of the (HACA)CON spectrum between 115 and 125 ppm of $^{15}N$ and 178 and 174 ppm of $^{13}C$ chemical shifts. **C:** Zoom-in on the proline region of the (HACA)CON spectrum.

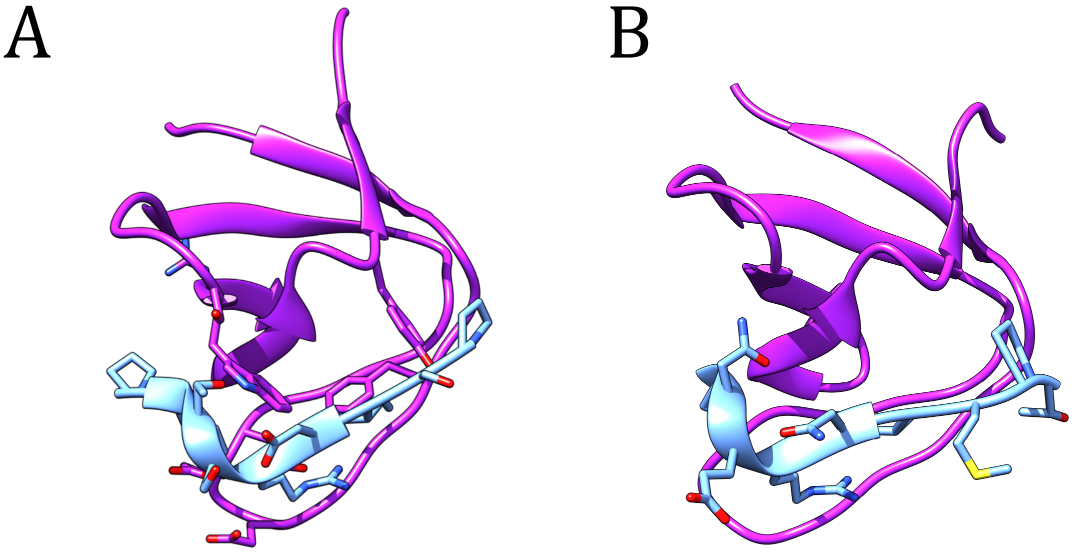


**Supplementary Figure S7:** Two complexes of SH3 domains and their respective Pro-rich peptides adopting a helical conformation. **A:** Complex between the GADS SH3 domain and a Pro-rich peptide derived from SLP76 (PDB: 2b0n). **B:** Complex between the STAM2 SH3 domain and a Pro-rich peptide derived from UBPY (PDB: 1uj0).

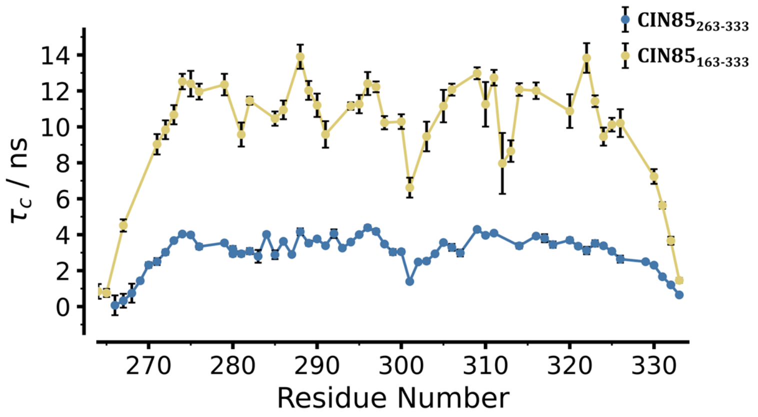


**Supplementary Figure S8:** Residue-specific apparent $\tau_c$ values determined using the TRACT experiment for CIN85$_{163–333}$ (yellow) and CIN85$_{263–333}$ (blue). The measurements were acquired at 800 MHz and a temperature of 298 K.

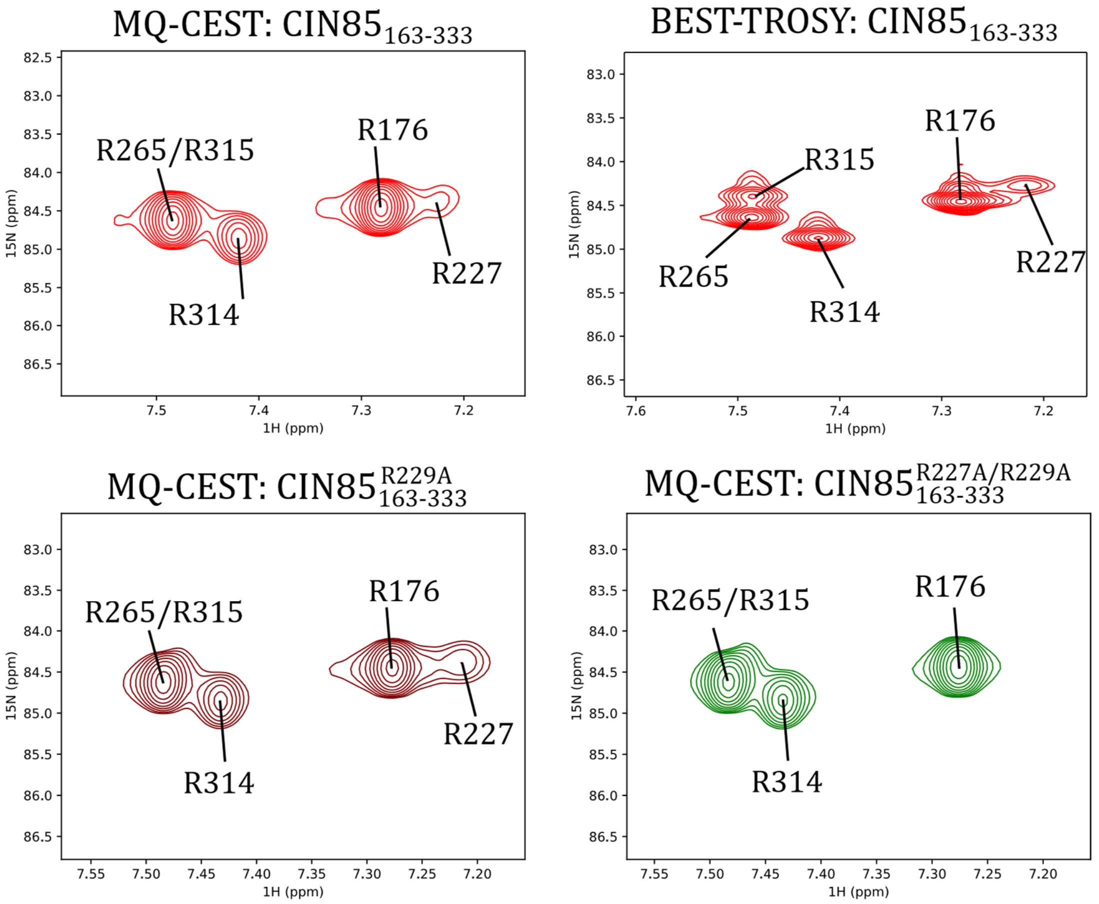


**Supplementary Figure S9:** Reference planes of the MQ-CEST experiments for the $CIN85_{163-333}$ constructs. For the WT construct, there is a comparison with the same spectral region of the BEST-TROSY to visualize the overlap of R265 and R315 in the MQ-CEST experiments.

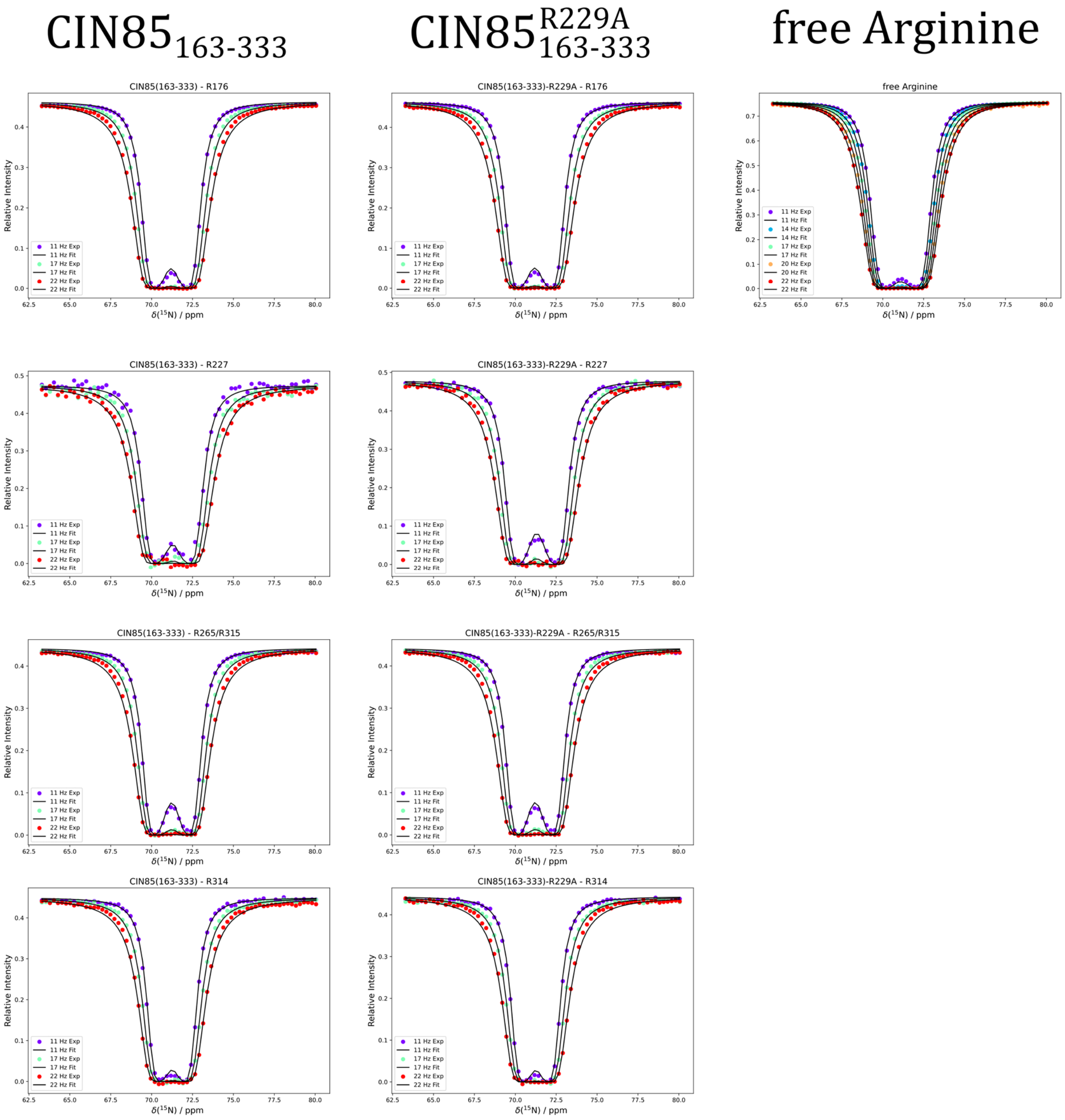


**Supplementary Figure S10:** MQ-CEST fits of the arginine sidechain peaks visible in the shorter CIN85 constructs (WT and R229A) at 800MHz. For comparison, the MQ-CEST profile of a $^{13}C/^{15}N$-labeled free arginine sample is also shown.

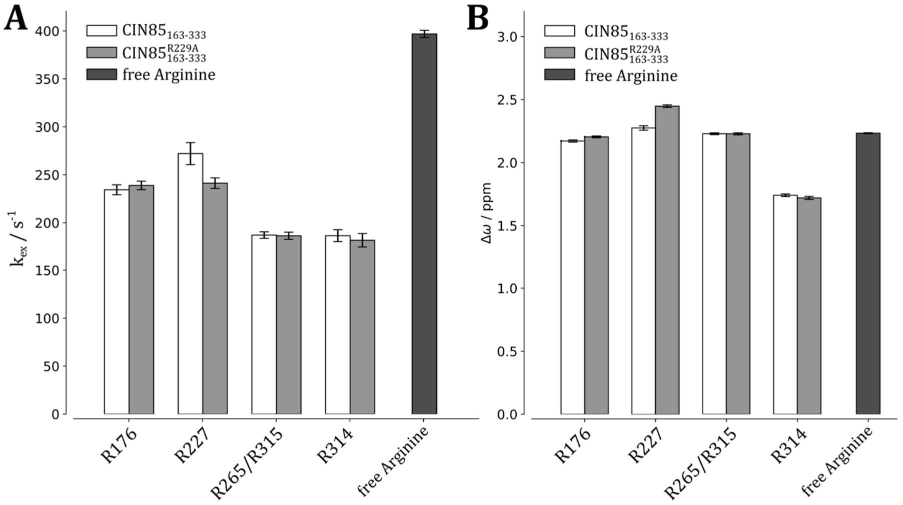


**Supplementary Figure S11:** Rates of rotation $k_{ex}$ (A) and chemical shift differences Δω between the two $N^{\eta}$ nuclei (B) of arginine sidechain guanidinium groups of uniformly $^{13}C/^{15}N$-labeled $CIN85_{163–333}$, $CIN85_{163–333}$-R229A and a sample of free arginine as determined via MQ-CEST experiments [16]. Experiments were carried out at 800 MHz and a temperature of 278K.

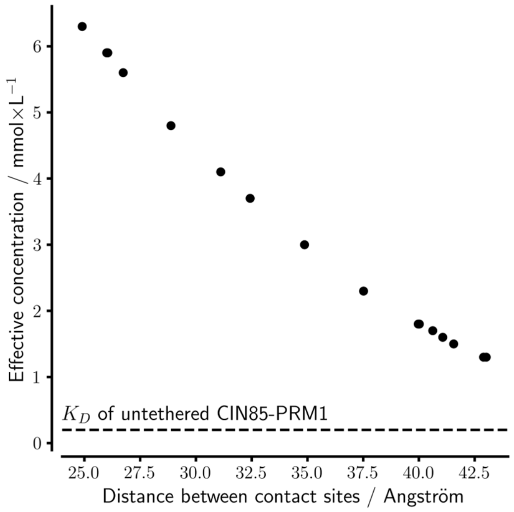


**Supplementary Figure S12:** Effective concentrations of the SH3C:CIN85-PRM1 complexes. These values were determined via the '$c_{eff}$ Calculator' web-server in dependence of the distance between contact sites. The distances between residues E232 and K269 used in the calculation were determined from the docked complex structures obtained using HADDOCK . The $K_D$ for the interaction of the untethered CIN85-PRM1 motif to SH3C is indicated as a dashed line (0.2 mM).

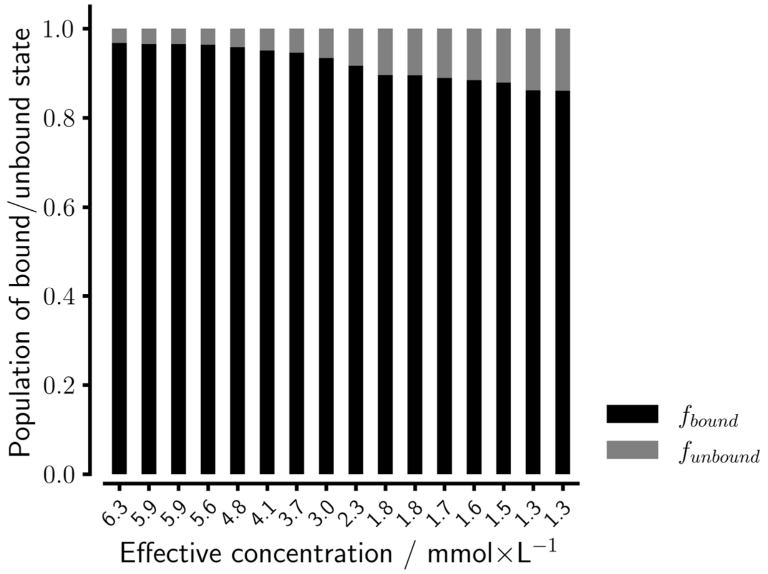


**Supplementary Figure S13:** Populations of the bound and unbound state of the SH3C:CIN85-PRM1 complex obtained from the effective concentrations. The effective concentrations used in the calculation were taken from Supplementary S12 and the corresponding $k_D$ from the titration of the untethered CIN85-PRM1 peptide to SH3C (see Table 1 in the manuscript).

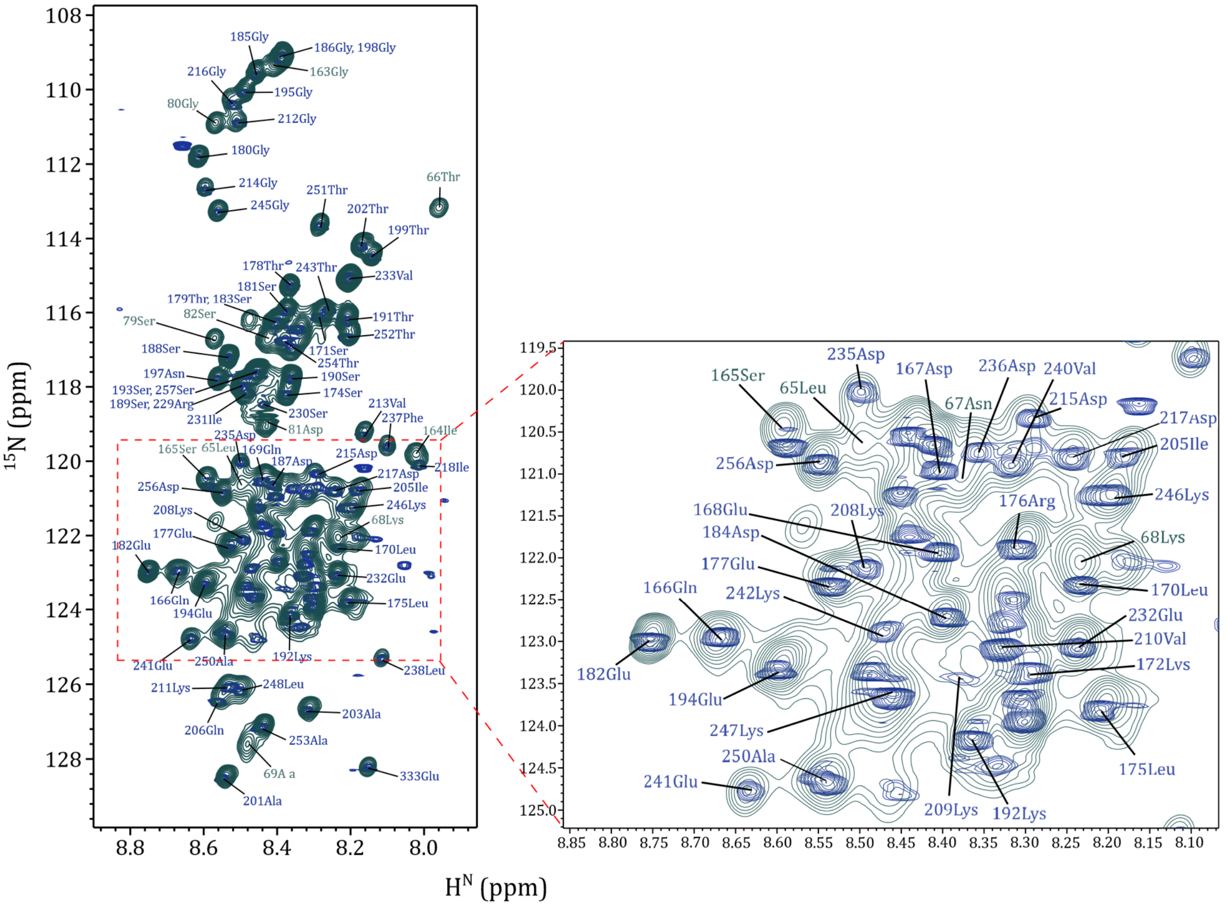


**Supplementary Figure S14:** Overlay of a $^{1}$H–$^{15}$N TROSY spectrum of 1mM uniformly $^{15}$N-$^{13}$C-labeled CIN85$_{1-333}$ at 1.2 GHz (green) and a $^{1}$H–$^{15}$N SOFAST-HMQC spectrum (blue) of 1 mM unlabeled CIN85$_{163-333}$ at 800MHz at natural abundance.

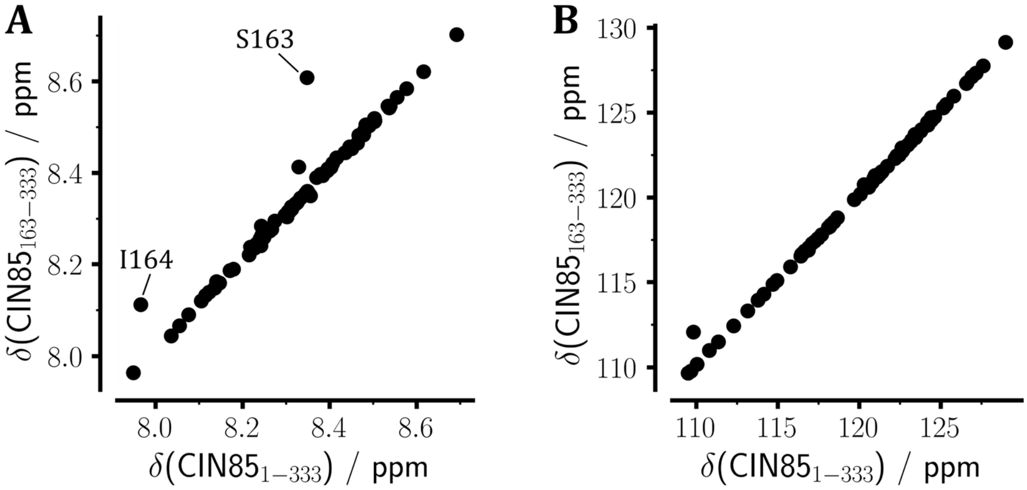


**Supplementary Figure S15:** Correlation plots of the assigned $^{1}H^{N}$ (A) and $^{15}N^{H}$ (B) chemical shifts of the flexible linker region in $CIN85_{163\text{-}333}$, compared to the assignment obtained using $CIN85_{1\text{-}333}$. Only the $^{1}H^{N}$chemical shifts at the N-terminal end of the construct (S163 and I164) exhibited significantly different chemical shifts from the longer $CIN85_{1\text{-}333}$ construct.

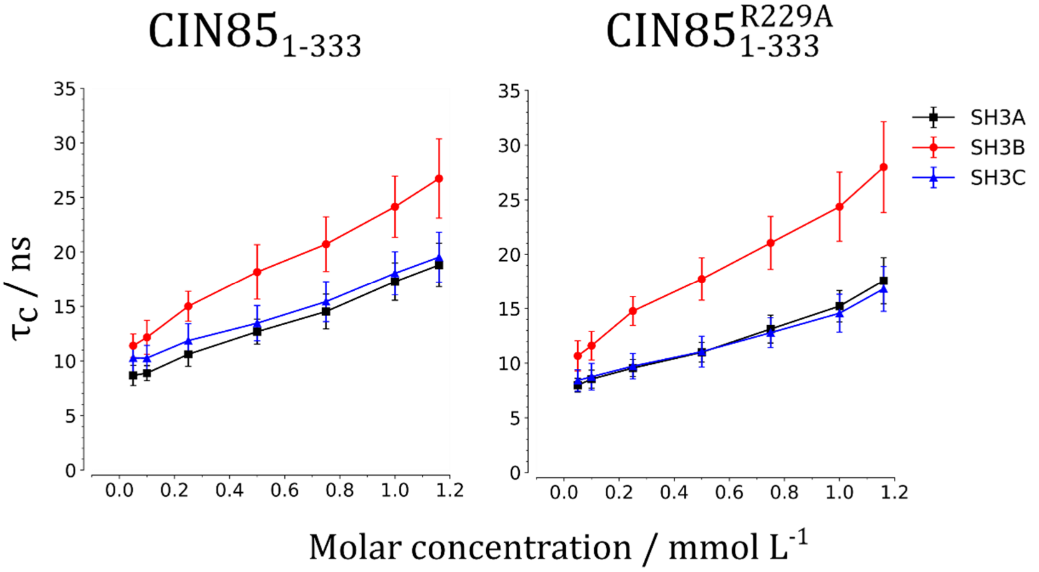


**Supplementary Figure S16:** Median rotational correlation time of the three SH3 domains of U[$^{2}H$,$^{15}N$]-labeled $CIN85_{1\text{–}333}$ (left) and $CIN85_{1\text{-}333}$-R229A (right) calculated from the residue-specific TRACT data, in dependence of the protein concentration. The error bars indicate the spread of the rotational correlation times throughout the SH3 domains, as calculated using the median absolute deviation.

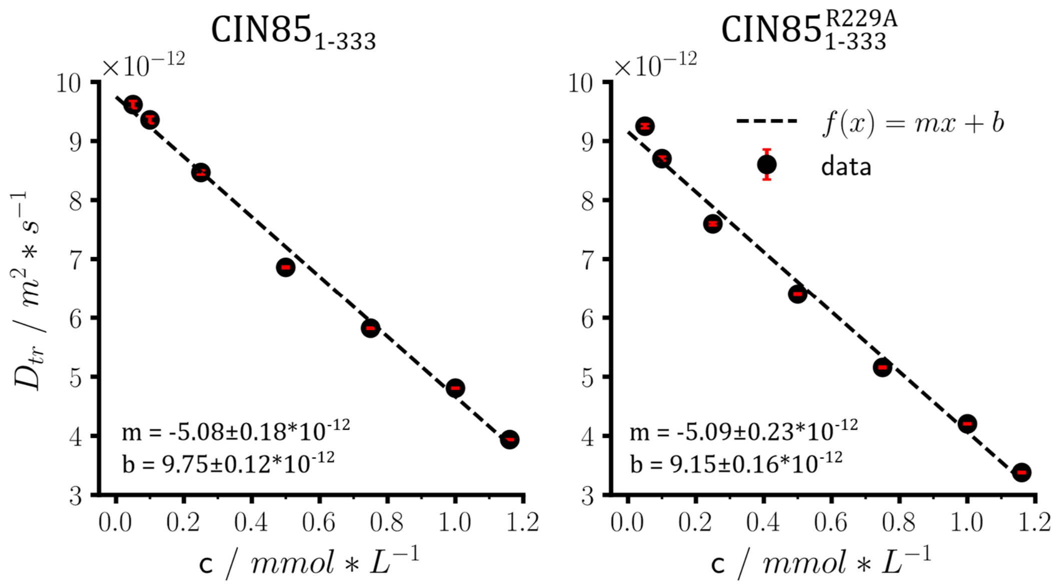


**Supplementary Figure S17:** Determination of the slope of the concentration-dependence of the diffusion coefficient of CIN85$_{1\text{–}333}$ and CIN85$_{1\text{–}333\text{-R229A}}$ via applying a linear fit of the form f(x) = mx + b. Here, m is the slope of the line and b its y-intercept.

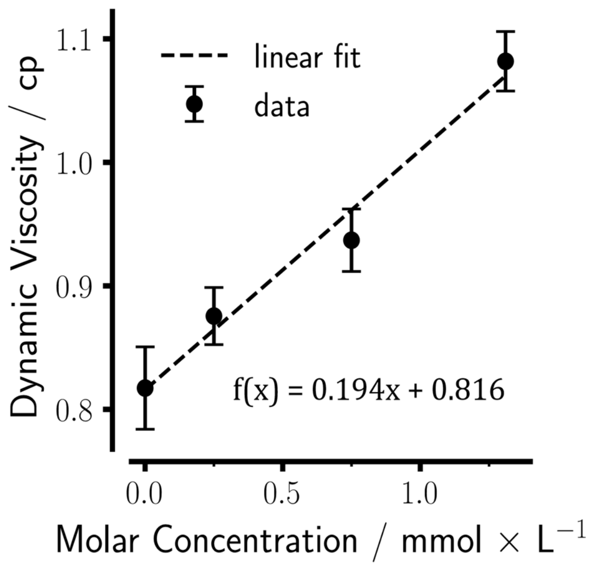


**Supplementary Figure S18:** Dynamic viscosity of U[$^{15}$N]-labeled CIN85$_{1\text{–}333}$ in the concentration range between 0 and 1.33 mmol L$^{-1}$. The zero point was just the measurement buffer (20 mM HEPES at pH 7.2, 100 mM NaCl, 1.0 mM TCEP, 0.5 mM Pefabloc, 0.5 mM EDTA, 0.01% $NaN_3$ and 5% $D_2O$). Viscosity was monitored via natural abundance $^{17}$O $T_1$ relaxation of water molecules as described elsewhere [22] at 298 K utilizing a Bruker Avance III HD 400MHz spectrometer.

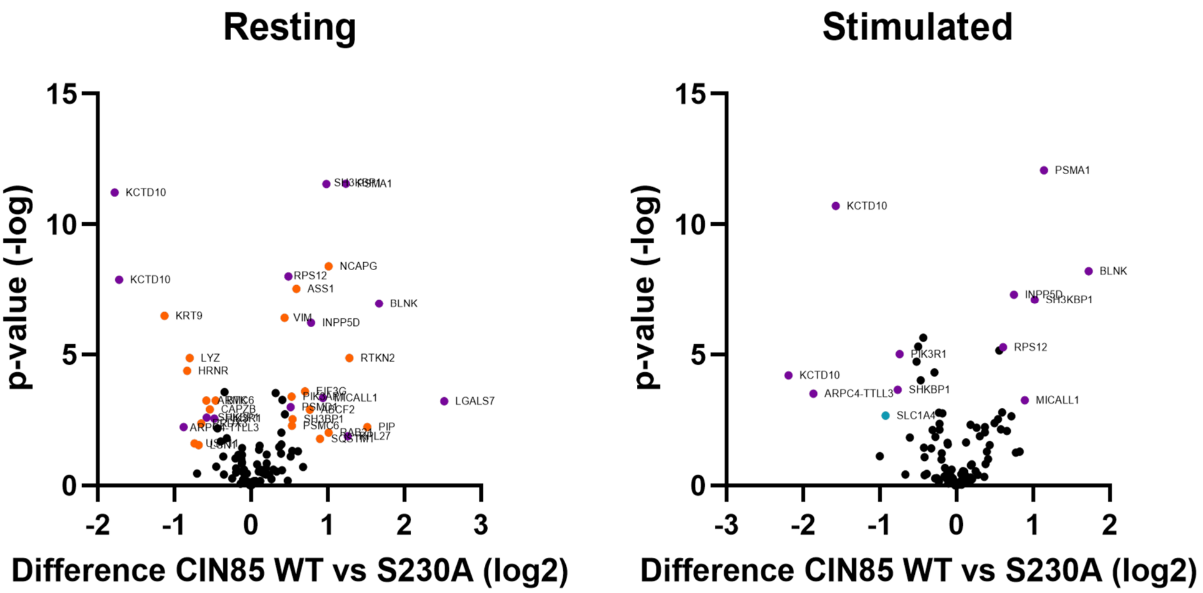


**Supplementary Figure S19:** Differences in the abundance of peptides purified from DG75 B cells carrying either CIN85 or CIN85-S230A in dependence of the activation state. Protein abundances were quantified using a label-free mass-spectrometric approach. The plots show the difference in the $log_2$ transformed abundances of proteins between citrine-CIN85 and citrine-CIN85-S230A for the resting (left) and stimulated state (right). Abundances of CIN85-S230A were subtracted from those of CIN85, yielding negative values in case of higher abundance compared to CIN85 and *vice versa*. The y-axis shows the associated p value and the proteins exhibiting statistically significant differences are labeled by their gene names. The color code is as follows: proteins that were statistically different between CIN85 and CIN85-S230A both in the resting and the stimulated state are colored purple, while the ones specific to one of the states are either orange (resting) or blue (stimulated).

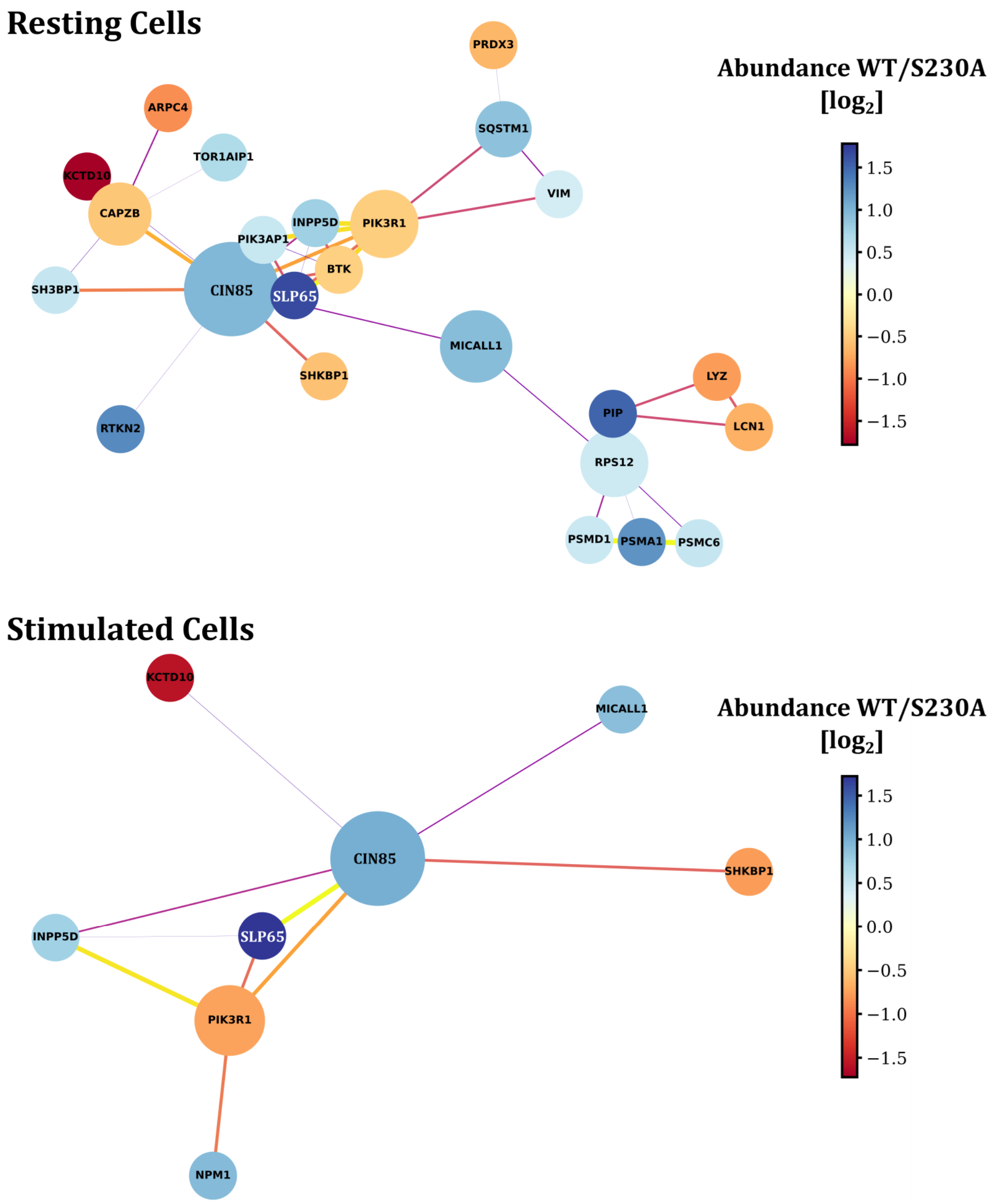


**Supplementary Figure S20:** STRING-based [23] network analysis based on the proteins that were found to have a significant difference in abundance in the mass-spectrometric experiments (see also Supplementary Table S3). Positive values (white-blue) indicate more abundance in citrine-CIN85 compared to the S230A mutant and negative values (orange-red) reflect a lower abundance in citrine-CIN85. The $log_2$-transformed values were normalized to show a symmetric range around zero for ease of visualization. The node-diameter reflects the interaction score from STRING.

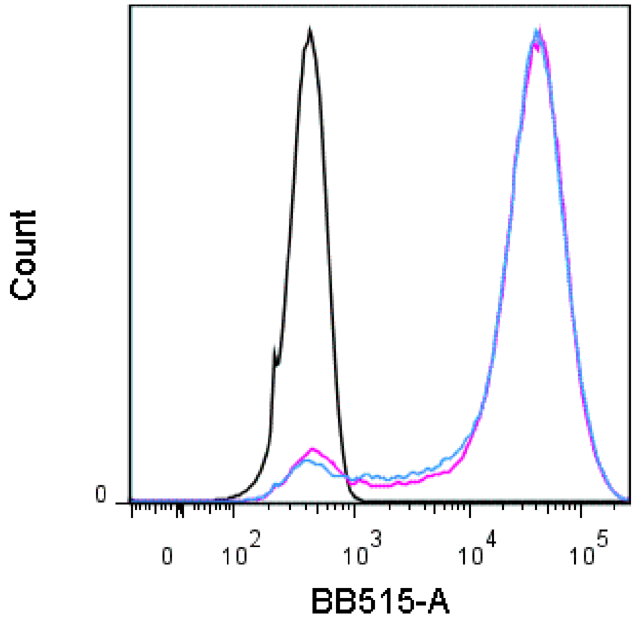


**Supplementary Figure S21:** Flow-cytometric determination of the expression level of citrine-labeled CIN85 (blue) and citrine-labeled CIN85-S230A (magenta) in DG75 B cells compared to the negative control (black).

### Supplementary Tables

#### Acquisition Parameters of the Resonance Assignment Experiments

**Supplementary Table S1**: Acquisition Parameters of $^{1}H$-detected triple resonance experiments used in the assignment of the CIN85 linker regions.

| Name | TD size($^{1}H/^{15}N/^{13}C$) | Scans | %NUS | Field / MHz |
|---|---|---|---|---|
| $CIN85_{1\text{-}333}$ | | | | |
| BEST-TROSY-HNCA | 2332/204/256 | 88 | 5 | 1200 |
| BEST-TROSY-HN(CO)CA | 2332/212/256 | 88 | 4 | 1200 |
| BEST-TROSY-HNCO | 1750/204/156 | 32 | 4 | 800 |
| $CIN85_{163\text{-}333}$ | | | | |
| BEST-TROSY-HNCACB | 1958/96/200 | 128 | 10 | 700 |
| BEST-TROSY-HN(CO)CACB | 1958/96/200 | 128 | 7.8 | 700 |

**Supplementary Table S2:** Acquisition Parameters of $^{13}C$-detected triple resonance experiments used in the assignment of the CIN85 linker regions.

| Name | TD size ($^{13}C/^{15}N/X$) | Scans | %NUS | Field / MHz |
|---|---|---|---|---|
| $CIN85_{163\text{-}333}$ | | | | |
| (HACA)N(CA)CON | 2048/80/108($^{15}N$) | 64 | 8.2 | 700 |
| (H)CCCON | 2048/80/128($^{13}C$) | 16 | 10 | 700 |
| H(CC)CON | 2048/44/120($^{1}H$) | 16 | 8 | 900 |
| C-HNCO | 1024/90/180($^{1}H$) | 64 | 10 | 800 |
| $CIN85_{163\text{-}333}$-R229A | | | | |
| (HACA)N(CA)CON | 2048/80/128($^{15}N$ | 80 | 8.1 | 700 |
| (H)CCCON | 2048/80/128($^{13}C$) | 16 | 8.5 | 700 |
| H(CC)CON | 2048/44/120($^{1}H$) | 16 | 8.5 | 700 |
| C-HNCO | 2048/84/112($^{1}H$) | 56 | 9.7 | 700 |
| $CIN85_{163\text{-}333}$-R227A/R229A | | | | |
| (HACA)N(CA)CON | 2048/80/118($^{15}N$ | 48 | 8.1 | 700 |
| (H)CCCON | 2070/80/128($^{13}C$) | 16 | 8.5 | 800 |

| | | | | |
|---|---|---|---|---|
| H(CC)CON | 2376/80/128($^1$H) | 16 | 8.5 | 800 |
| C-HNCO | 2048/84/112($^1$H) | 32 | 9.7 | 700 |

### Protein abundances determined via label-free Mass Spectrometry

**Supplementary Table S3:** Abundances of peptides found in the label-free mass-spectrometric analysis of DG75 B cells. The values shown were calculated as differences in the abundance of peptides in the CIN85-S230A mutant DG75 B cells compared to cells carrying the wild-type CIN85. Negative values indicate an increased interaction to the S230A mutant while proteins with positive values were found to associate less with CIN85-S230A.

| | | CIN85/CIN85-S230A | |
|---|---|---|---|
| Protein | **Uniprot Entry Name** | **Basal** | **Stimulated** |
| Actin Regulation | | | |
| CAPZB | CAPZB_HUMAN | -0.541 | / |
| ARPC4 | ARPC4_HUMAN | -0.881 | -1.868 |
| PIP | PIP_HUMAN | 1.515 | / |
| Proteasome | | | |
| PSMA1 | PSMA1 | 1.235 | 1.142 |
| PSMD1 | PSMD1_HUMAN | 0.514 | / |
| PRS10 | PRS10_HUMAN | 0.533 | / |
| Scaffold Proteins | | | |
| BLNK (SLP65) | BLNK_HUMAN | 1.668 | 1.722 |
| SH3KBP1 (CIN85) | SH3KBP1_HUMAN | 0.983 | 1.017 |
| SHKBP1 | SHKBP1_HUMAN | -0.581 | -0.769 |
| PI3K Signaling | | | |
| SHIP-1 | SHIP1_HUMAN | 0.783 | 0.747 |

| | | | |
|---|---|---|---|
| BCAP | BCAP_HUMAN | 0.528 | / |
| P85A | P85A_HUMAN | -0.48 | -0.743 |
| Mitochondrial | | | |
| HIG1A | HIGD1A_HUMAN | / | -2.678 |
| AT5F1 | AT5F1_HUMAN | / | 1.13 |
| MTDC | MTDC_HUMAN | / | -2.857 |
| PRDX3 | PRDX3_HUMAN | -0.652 | / |
| Rho GTPase Regulation | | | |
| 3BP1 | 3BP1_HUMAN | 0.54 | / |
| Ubiquitination | | | |
| KCTD10 | BACD3_HUMAN | -1.722 | -2.193 |
| UBP11 | UBP11_HUMAN | -0.74 | / |
| Chromatin | | | |
| CND3 | CND3_HUMAN | 1.011 | / |
| Ribosome | | | |
| RS12 | RS12_HUMAN | 0.483 | / |
| Filaments | | | |
| VIME | VIME_HUMAN | 0.434 | / |
| Nucleus | | | |
| TOIP1 | TOIP1_HUMAN | 0.677 | / |
| COIL | COIL_HUMAN | 0.635 | / |
| NPM | NPM_HUMAN | / | 0.926 |
| B cell Development | | | |
| RTKN2 | RTKN2_HUMAN | 1.281 | / |

| | | | |
|---|---|---|---|
| Btk | BTK_HUMAN | -0.466 | / |
| Trafficking | | | |
| OSBL8 | OSBL8_HUMAN | 1.174 | / |
| MILK1 | MILK1_HUMAN | 0.937 | 0.89 |
| RAB21 | RAB21_HUMAN | 1.008 | / |
| ENPL | ENPL_HUMAN | / | 0.646 |
| Autophagocytosis | | | |
| SQSTM | SQSTM_HUMAN | 0.898 | / |
| Miscellaneous | | | |
| LYSC | LYSC_HUMAN | -0.803 | / |
| ASSY | ASSY_HUMAN | 0.588 | / |
| HORN | HORN_HUMAN | -0.835 | / |
| ARMC6 | ARMC6_HUMAN | -0.586 | / |
| LEG7 | LEG7_HUMAN | 2.519 | / |
| ABCF2 | ABCF2_HUMAN | 0.763 | / |
| LCN1 | LCN1_HUMAN | -0.684 | / |
| KCY | KCY_HUMAN | / | 0.755 |
| SATT | SATT_HUMAN | / | -0.929 |

**Experiment Definitions of $^{13}$C-Detected Resonance Assignment Experiments for the Semi-Automatic Assignment using FLYA**

```
SPECTRUM hacaCON CO N
  0.980 CO:C_BYL (C_ALI N_AMI) N:N_AMI

SPECTRUM hacaNcaCON N_1 N_2 CO
  0.980 CO:C_BYL N_1:N_AMI C_ALI C_BYL
  0.980 N_1:N_AMI C_ALI CO:C_BYL N_2:N_AMI

SPECTRUM hCCCON CO N C
  0.98 N:N_AMI (C_ALI) CO:C_BYL C:C_ALI
  0.98 N:N_AMI (C_ALI) CO:C_BYL C_ALI C:C_ALI
  0.80 N:N_AMI (C_ALI) CO:C_BYL C_ALI C_ALI C:C_ALI
  0.60 N:N_AMI (C_ALI) CO:C_BYL C_ALI C_ALI C_ALI C:C_ALI
  0.40 N:N_AMI (C_ALI) CO:C_BYL C_ALI C_ALI C_ALI C_ALI C:C_ALI

SPECTRUM HccCON CO N H
  0.98 N:N_AMI (C_ALI) CO:C_BYL C_ALI H:H_ALI
  0.98 N:N_AMI (C_ALI) CO:C_BYL C_ALI C_ALI H:H_ALI
  0.80 N:N_AMI (C_ALI) CO:C_BYL C_ALI C_ALI C_ALI H:H_ALI
  0.60 N:N_AMI (C_ALI) CO:C_BYL C_ALI C_ALI C_ALI C_ALI H:H_ALI
  0.40 N:N_AMI (C_ALI) CO:C_BYL C_ALI C_ALI C_ALI C_ALI C_ALI H:H_ALI

SPECTRUM CBCACON CO N C
  0.98 N:N_AMI (C_ALI) CO:C_BYL C:C_ALI
  0.98 N:N_AMI (C_ALI) CO:C_BYL C_ALI C:C_ALI

SPECTRUM CONH CO N H
0.98 CO:C_BYL (C_ALI N_AMI) N:N_AMI H:H_AMI
```
